# The oxygen-induced thiosulfate production during Sulfate reduction, Autotrophic denitrification, Nitrification and Anammox (SANIA) integrated process towards next-generation mainstream wastewater treatment

**DOI:** 10.1101/2023.06.12.544556

**Authors:** Chu-Kuan Jiang, Yang-Fan Deng, Guang-Hao Chen, Di Wu

## Abstract

This study proposes a novel integrated process: the oxygen-induced thiosulfate production during <u>S</u>ulfate reduction, <u>A</u>utotrophic denitrification, <u>NI</u>trification and <u>A</u>nammox (SANIA) integrated process, targeting to treat mainstream wastewater after organics capture. Three moving-bed biofilm reactors (MBBRs) were applied for oxygen-induced thiosulfate production during sulfate reduction, mixed sulfide- and thiosulfate-driven partial denitrification and anammox (MSPDA), and nitrification (N), respectively. This study firstly established SANIA with supply of mimic nitrifying effluent to investigated the development of MSPDA (Period I) and subsequently verified SANIA in mainstream condition with implementation of nitrification (Period II). In Period I, the MSPDA-MBBR fed with ERATO-MBBR and mimic nitrifying effluent, was operated for over 300 days. Without inoculation of anammox biomass, the high rates of denitratation and anammox being 2.7 gN/(m^2^·d) and 2.8 gN/(m^2^·d), respectively were developed in the bioreactor with the enrichment of anammox bacteria and the coexistence of sulfur-oxidizing bacteria. Batch tests were performed to explore the bioconversion of sulfur and nitrogen compounds in MSPDA with main findings as following: 1) kinetics and pathways of sulfide (S^2-^ ➔ S^0^ ➔ SO_4_^2-^) and thiosulfate (S_2_ O_3_^2-^ ➔ SO_4_^2-^) oxidation were revealed; 2) fast denitrification was achieved during oxidation of sulfide and thiosulfate to S^0^ and sulfate, respectively with sufficient nitrite accumulation, supporting the high activity of anammox; 3) nitrite utilization rate of anammox (50.8 mgN/(m^2^·h)) is higher than sulfur-driven denitritation (12.9−42.6 mgN/(m^2^·h)), demonstrating the dominance of anammox in nitrogen removal. In Period II, the N-MBBR was set behind MSPDA-MBBR to supply nitrate by recirculation, thus the SANIA system was developed. Afterwards the integrated SANIA system with a short HRT of 4.7 h was continuously operated for over 130 days. Results demonstrated that 90% of COD, 93% of ammonium and 61% of TIN were removed with concentration of COD, ammonium, and TIN below 10 mg/L, 3 mgN/L and 13 mgN/L, respectively in effluent. Combining with organic capture and SANIA for sewage treatment, the energy-neutral and space-efficient treatment of mainstream wastewater is promising.

**Highlights:**

- A new sulfur-cycle process (SANIA) was developed for sewage treatment.
- High rates of both denitratation and anammox were developed in MSPDA.
- Oxidation of TdS (to S^0^) and S_2_O_3_^2-^ (to SO_4_^2-^) in MSPDA boosts nitrite buildup.
- 74−81% of the removed TIN was via anammox in SANIA process.
- SANIA with a short HRT of 4.7h achieved good effluent qualities for sewage treatment.

## 1. Introduction

The annual electricity input for WWTPs accounts for about 3% of the electricity load in the world (<u>IEA, 2022</u>). Aeration energy could contribute to 45−70% energy consumption of WWTP for organics and nitrogen removal in conventional activated sludge process (Rosso et al., 2008). The application of anaerobic ammonium oxidation (anammox) process, requiring low aeration energy and no organics for nitrogen removal, could maximumly reducing 60% of aeration energy and recovering 90% of organics as energy source through combination of organics harvesting processes (Harleman and Murcott, 1999; Jimenez et al., 2015; McCarty et al., 2011). The minimization of energy input and maximization of organics recovery could make mainstream wastewater treatment energy-neutral (Kartal et al., 2010). However, the realization of anammox in mainstream is still hardly to be demonstrated over a decade of efforts (Cao et al., 2017; Wang et al., 2022).

Partial nitrification anammox (PN/A) and heterotrophic partial denitrification anammox (PD/A) have been intensively studied recently for demonstration of anammox in mainstream wastewater treatment, yet difficult to be applied due to complex control for stable nitrite provision (e.g., residual ammonium control or pH control via introducing anaerobic digestion liquor) or heterotrophic biomass (1.1 gVSS/gNO_3_^-^-N) overgrowth (Cao et al., 2017; Du et al., 2016; Henze et al., 2006; Wang et al., 2021). Notably in the emerging sulfur-based autotrophic PD/A (SPDA) process (Deng et al., 2022), reduced sulfur compounds (sulfide, thiosulfate and S^0^) were demonstrated to be effective electron donors for the stable partial denitrification (NO_3_^-^➔NO_2_^-^), then, produced nitrite could be utilized with ammonium for anammox (NH_4_^+^+NO_2_^-^➔N_2_). Besides, low risk of outcompeting anammox biomass is another advantage of SPDA due to low yield (0.15–0.57 g VSS/gNO_3_^-^-N) of sulfur oxidizing bacteria (SOB) (Cui et al., 2019a). Thus, SPDA is a promising alternative for applying anammox in mainstream wastewater treatment.

However, the SPDA developed in wastewater treatment still relies on the external dosage of reduced sulfur compounds, which is not economical enough in practice (Deng et al., 2022). A integrated process, like conventional activated sludge, sulfate reduction, autotrophic denitrification, nitrification integrated (SANI^®^), PN/A, or heterotrophic PD/A process without external dosage of chemicals (Gilbert et al., 2014; Ludzack and Ettinger, 1962; Ma et al., 2017; Wang et al., 2009a), for treating mainstream wastewater is yet to be explored. As oxygen-induced thiosulfate production during sulfate reduction has been developed with main products of both sulfide and thiosulfate for treatment of mainstream wastewater commonly containing sulfate of 20−180 mgS/L after harvesting organics due to seawater intrusion, application of seawater toilet flushing, and dosage of sulfate-based coagulants (Jiang et al., in press), here we propose the oxygen-induced thiosulfate production during Sulfate reduction, Autotrophic denitrification, NItrification and Anammox (SANIA) integrated process for achieving mainstream anammox. In the SANIA process as shown in Fig. 1: organic are firstly utilized in ERATO with sulfide and thiosulfate produced as main products with the effect of oxygen; and then the mixed sulfide- and thiosulfate-driven SPDA (MSPDA) could simultaneously and fast remove ammonium and nitrate, the ammonium and nitrate are supplied by ERATO effluent and recycled nitrifying effluent respectively; at last, the residual ammonium in the effluent of MSPDA is oxidized to nitrate through nitrification. This process will theoretically need zero dose of electron donors to energy-efficiently treat mainstream wastewater.

**Fig. 1.**
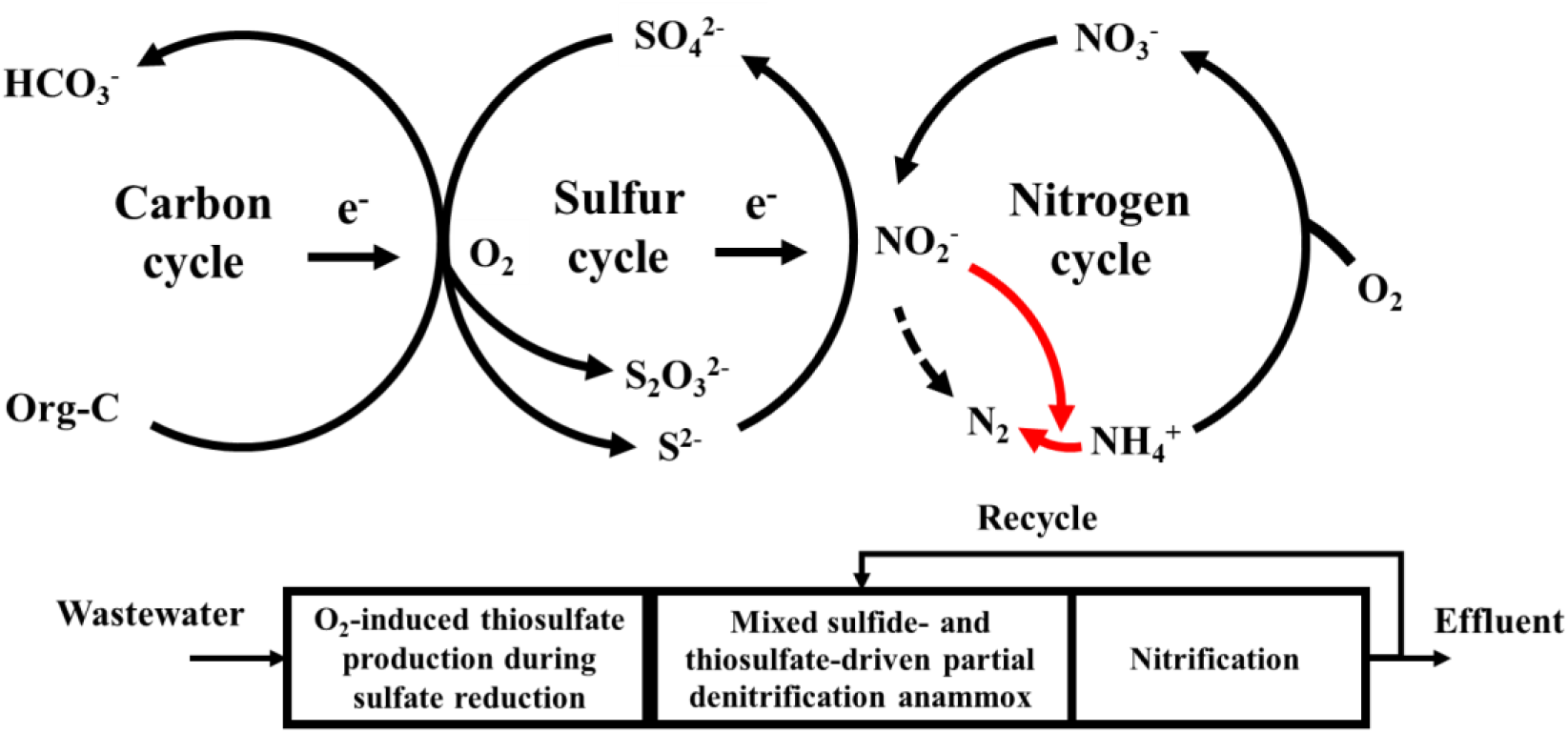
The conceptional SANIA process operated in three reactors (the red line refers to anammox pathway).

Nevertheless, the development of SANIA faces several uncertainties and challenges for treating mainstream wastewater containing low nitrogen of 20−75 mgN/L (Deng et al., 2022; Volcke et al., 2020): (1) nitrite provision for anammox might not necessarily be achieved as Moraes et al., (2012) reported that nitrite was preferred to be reduced by SOB under a sulfide to nitrate (S/N, unit: gS/gN) ratio of 1.1−1.9; (2) anammox biomass was inevitably inoculated in previous SPDA processes (Deng et al., 2019; Deng et al., 2021a; Yang et al., 2020), the direct enrichment of anammox biomass in MSPDA without inoculation is questionable; (3) as the rates of sulfur-driven denitrification and anammox are as high as 0.4 gN/(L·d) and 0.3−0.6 gN/(L·d), respectively (Cui et al., 2019b; Laureni et al., 2016), a practical nitrogen removal rate much higher than 0.1 gN/(L·d) obtained in previously SPDA might be achieved (Yang et al., 2020); (4) the nitrogen removal efficiency of SANIA might be affected the residual dissolved oxygen in nitrifying effluent due to consumption of limited sulfide of 14−20 mgS/L and thiosulfate of 5−10 mgS/L produced in sulfate-reducing effluent (Jiang et al., in press).

To clarify the uncertainties mentioned above, the feasibility of MSPDA was investigated first in the SANIA system with mimic nitrifying effluent applied, aiming to reveal the insights of MSPDA including the development of denitrification and anammox, the enrichment of responsible bacteria, and the sulfur (S) and nitrogen (N) compounds conversion. After which, the performance of SANIA system was further investigated and evaluated with integration of nitrification for nitrate supply. Through establishment of SANIA, this study aims to provide a promising solution for sustainable wastewater treatment.

## 2. Material and methods

### 2.1 Reactor set-up and operation

For establishment of SANIA system, the operation was divided into two periods with mimic and real nitrifying effluent applied, respectively as shown in Supplementary Information (SI) 1. In Period I (days 1–319), two moving-bed biofilm reactors (namely ERATO-MBBR and MSPDA-MBBR) were installed and continuously operated. Among which, ERATO- MBBR fed with synthetic mainstream wastewater was applied as reported (Jiang et al., in press). Effluent of which mixed with nitrate, containing fluctuated sulfide (10–16 mgS/L), thiosulfate (4–8 mgS/L), ammonium (28–30 mgN/L), and nitrate (30 mgN/L) with NO_3_^-^-N/NH_4_^+^-N as approximately 1:1, was applied as influent for MSPDA-MBBR. The hydraulic retention time (HRT) and working volume for both ERATO-MBBR and MSPDA-MBBR were 0.7 h and 1 L, respectively. In Period II (days 325–455), the effluent of MSPDA-MBBR was fed into a third reactor of N-MBBR and the effluent of which was recirculated back to MSPDA-MBBR for nitrate supply. The HRT of MSPDA-MBBR was extended to 1.1 h to fully utilize the electron donors in effluent of ERATO-MBBR by applying a working volume of 1.6 L. Nominal HRTs of ERATO-MBBR and N-MBBR were 0.7 h and 3 h, resulting in total HRT of 4.7 h for SANIA. No sludge was purposely wasted from the system.

MSPDA-MBBR and N-MBBR were inoculated with K3 AnoxKaldnes^TM^ carriers and ActiveCell 920 carriers taken separately from the parent reactors (as described in SI 1) operated in our lab previously. The carrier filling ratios of MSPDA-MBBR and N-MBBR were 50% and 45%, respectively. A magnetic stirrer with the stirring speed of 160 rpm and the aeration with flow rate of 2.5 L/min were applied in MSPDA-MBBR and N-MBBR, respectively. Temperature of MSPDA-MBBR and N-MBBR were maintained at 25 ℃. pH in two reactors varied at 7–8 without intentional control as summarized in Table 1.

**Table 1.**
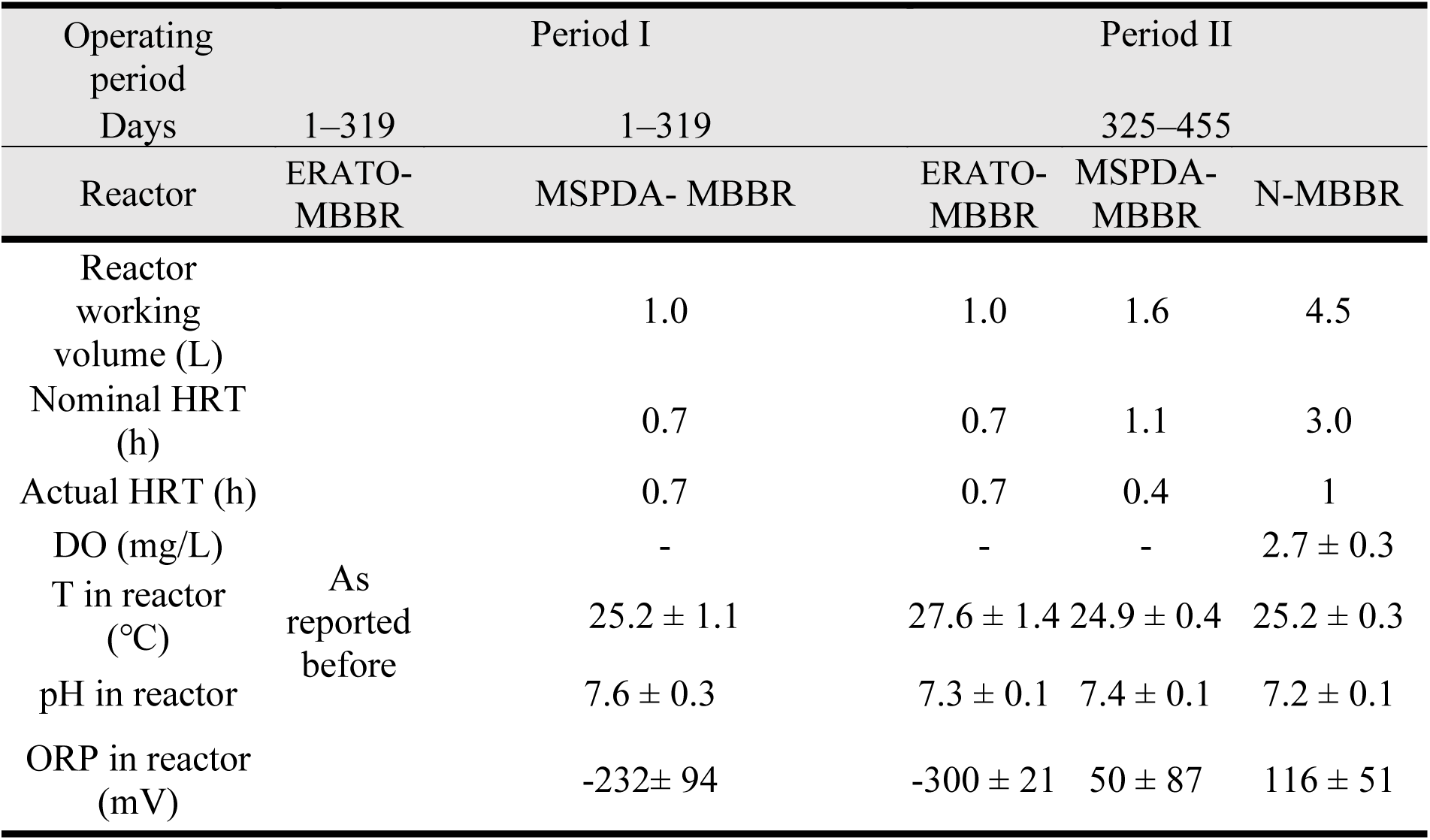
Operation conditions in Period I (Days 1–319) and Period II (Days 325–455).

### 2.2 Microbial community analysis

Microbial communities were analyzed in both Period I and Period II. In Period I, the biofilm samples were taken from MSPDA-MBBR on day 106, 240 and 319. After implement of N-MBBR in Period II, another 3 biofilm samples from ERATO-/MSPDA-/N-MBBR were taken on day 455. For microbial community analysis, the universal forward primer 515F (5’- GTGCCAGCMGCCGCGG-3’) and the reverse primer 907R (5’- CCGTCAATTCMTTTRAGTTT-3’) were used for amplifying V4 and V5 regions of the 16S rRNA gene (Quince et al., 2011). At least 60 ng DNA was extracted from each sample for 16S rDNA gene PCR amplification which was conducted on Illumina PE250 platform (Novogene, Hong Kong). The raw tags were dealt with quality filtering through Qiime (V1.7.0) for obtaining clean tags. Based on which, OTU clustering was then performed with 97% identity by using Uparse software. At last, species annotation was performed by Qiime (Version 1.7.0) with comparing SSUrRNA database of SILVA Database. The raw data of the 6 samples in total were uploaded in National Center for Biotechnology Information (NCBI) with a BioProject ID of PRJNA966788.

### 2.3 Batch tests

Four batch tests (A, B, C, D) were conducted after MSPDA-MBBR operated 319 days to investigate i) the kinetic rates of sulfide- and thiosulfate-driven denitrification, and anammox (batch A, B, and C); and ii) the S and N compounds conversion in MSPDA (batch D).

Batch A and B were conducted to specifically investigate kinetic rates of sulfide and thiosulfate oxidation. TdS (25 mgS/L) and thiosulfate (50 mgS/L) were employed as the electron donor in batch A and B, respectively, and 30 mg N/L of nitrate served as electron acceptor in both tests (Table 2). And no ammonium was provided to restrict anammox activity. The batch C was performed with 30 mg N/L of ammonium and 30 mg N/L of nitrite to investigate the anammox activity of biofilm cultivated in MSPDA-MBBR. Finally, batch D studied the overall S and N conversion in the MSPDA, in which 15 mg S/L TdS, 20 mgS/L thiosulfate, 30 mg N/L ammonium and 30 mg N/L nitrate were set. The samples with 2−4 mL volume were taken every 15−45 min to analyze sulfate, sulfide, thiosulfate, ammonium, nitrite, and nitrate. All the tests were performed in duplicate. In each batch test, five carriers from the MSPDA-MBBR were collected and washed with deoxygenated ultrapure water for three times. The specific surface area was 0.1 m^2^/L since batch bottle applied for batch test had a working volume of 205 mL. Then, carriers and 190 mL deoxygenated ultrapure water were added to batch bottles, and nitrogen gas was sparged for 30 min to remove dissolved oxygen (DO) from headspace and liquid phase. At last, total 10−20 mL of stock solutions (including sulfide, thiosulfate, ammonium, nitrite, nitrate, alkalinity, and etc.) were injected to batch bottles to achieve the designed initial concentrations. Experimental temperature was controlled at 30 °C via water bath shaker with shaking rate of 180 rpm. 0.1M H_3_PO_4_ and 0.1 M NaOH were dosed to adjust the initial pH at around 7.5.

**Table 2.** Details of batch conditions.

| Batch | COD<sub>sulfur</sub>*<br>(mg/L) | TdS<br>(mgS/L) | S<sub>2</sub>O<sub>3</sub><sup>2-</sup><br>(mgS/L) | NH<sub>4</sub><sup>+</sup><br>(mgN/L) | NO<sub>2</sub><sup>-</sup><br>(mgN/L) | NO<sub>3</sub><sup>-</sup><br>(mgN/L) |
| --- | --- | --- | --- | --- | --- | --- |
| A | 50 | 25 | - | - | - | 30 |
| B | 50 |  | 50 | - | - | 30 |
| C | - | - | - | 30 | 30 | - |
| D | 50 | 15 | 20 | 30 | - | 30 |

COD<sub>sulfur</sub>\*: indicates the equivalent COD of reduced sulfur compounds in perspective of contained electrons.

### 2.4 Calculations

For nitrifying effluent recirculation in SANIA system, the recycle ratio of 2.0 was determined through a detailed calculation conducted in SI 2.1 to achieve a TIN of lower than 10 mg/L, according to the theoretical steady-state operation of SANIA. For analysis of the data obtained from both SANIA operation and batch tests, the kinetic rates including the surface-specific rates of sulfur-driven denitrification, anammox, and nitrification were calculated as provided in SI 2.2. The contributions of anammox and denitrification to TIN removal were evaluated as shown in SI 2.3. The electron balance was calculated as accepted electrons for both denitratation and denitritation divided donated electrons from oxidation of both sulfur compounds and organics, and the sulfur loss (S^0^) and nitrate production by anammox were considered in this calculation. The mass balance for sulfur was calculated as (ΔS_2_O_3_^2−^-S + ΔTdS-S)/|ΔSO_4_^2−^-S| × 100%.

### 2.5 Chemical analysis

During the experiments, the influent and effluent samples of all reactors were collected 2−4 times a week. All samples were filtered with 0.22 μm Millipore^TM^ syringe filters before chemical analysis. TdS was measured immediately using methylene blue method (APHA, 2005). Ammonium, nitrite and nitrate was determined by spectrophotometric method (APHA, 2005). Sulfate and thiosulfate were measured with an ion chromatographer HIC-20A super (Shimadzu Corp, Japan) equipped with an IC-SA2 analytical column. Total dissolved organic carbon (TOC) was analyzed with a Total Organic Carbon Analyzer (Shimadzu TOC-5000A). The conversion factor between TOC to COD was determined to be 2.66 via measurements. Alkalinity, suspended solid (SS), and volatile suspended solid (VSS) in bulk liquid were measured according to the Standard Method (APHA, 2005). The attached total solids (ATS, mg ATS·L^-1^) and attached volatile solids (AVS, mg AVS· L^-1^) of the MBBR biofilm were measured according to Cui et al., (2019c). The effluent of reactors was also collected for the measurement of DO concentration, pH and temperature by a multiparameter meter YSI Pro Plus (YSI, USA).

## 3. Results and discussion

### 3.1 SANIA in Period I: investigation of MSPDA-MBBR

#### 3.1.1 Development of MSPDA in continuous reactors

Lone-term performance of MSPDA-MBBR in Period I was shown in Fig. 2. The MSPDA-MBBR in was continuously operated for 319 days, fed with the mixture of sulfate-reducing effluent and mimic nitrifying effluent containing TdS, thiosulfate, ammonium and nitrate with concentrations of 12.1–16.0 mgS/L, 3.9–10.4 mgS/L, 28.5–29.9 mgN/L, and 29.3–34.7 mgN/L respectively as shown in SI 3. With sufficient nitrate supply, both TdS and thiosulfate were fully utilized in MSPDA-MBBR throughout the period. Meanwhile, the concentration of nitrite in effluent was 5.8–6.9 mgN/L, indicating the effectiveness of nitrite provision. For the development of MSPDA, the start-up of autotrophic denitrification was indicated by surface-specific removal rates of TdS, thiosulfate and nitrate, reached 1.7 gS/(m^2^·d), 1.5 gS/(m^2^·d) and 2.7 gN/(m^2^·d), respectively during days 98–135. Meanwhile, the ammonium removal rate of 0.7 gN/(m^2^·d) indicated the emergence of anammox. After which in days 136–319, the removal rates of TdS, thiosulfate, and nitrate achieved and maintained at 2.3 gS/(m^2^·d), 0.6–0.9 gS/(m^2^·d), and 2.7–2.9 gN/(m^2^·d) respectively, indicating the stabilization of autotrophic denitrification. The obtained nitrate removal rate was much higher than it of 1.1 gN/(m^2^·d) in previous autotrophic denitrification MBBR (Cui et al., 2019c). Specifically, the nitrate removal rate was increased from 2.7 gN/(m^2^·d) to 2.9 gN/(m^2^·d) when the thiosulfate rose from 3.9 ± 2.4 mgS/L to 6.4 ± 2.9 mgS/L on day 241, indicating that the increase of thiosulfate is beneficial to fast sulfur-driven denitrification as reported (Cardoso et al., 2006). Meanwhile, the ammonium removal rate achieved and maintained at 1.4 gN/(m^2^·d) from day 241 on, indicating that anammox performance became stable at last in MSPDA. In this MSPDA-MBBR, it should be noted that that organics removal of 11.5 mg COD/L was detected in the bioreactor after performance stabilized, suggesting that sulfur-driven autotrophic denitrification and heterotrophic denitrification contributed to 80% and 20% of the nitrate removal, respectively. The calculation is based on the electron donation in oxidation of sulfur compounds and organics with electron balance of 88%.

**Fig. 2.**
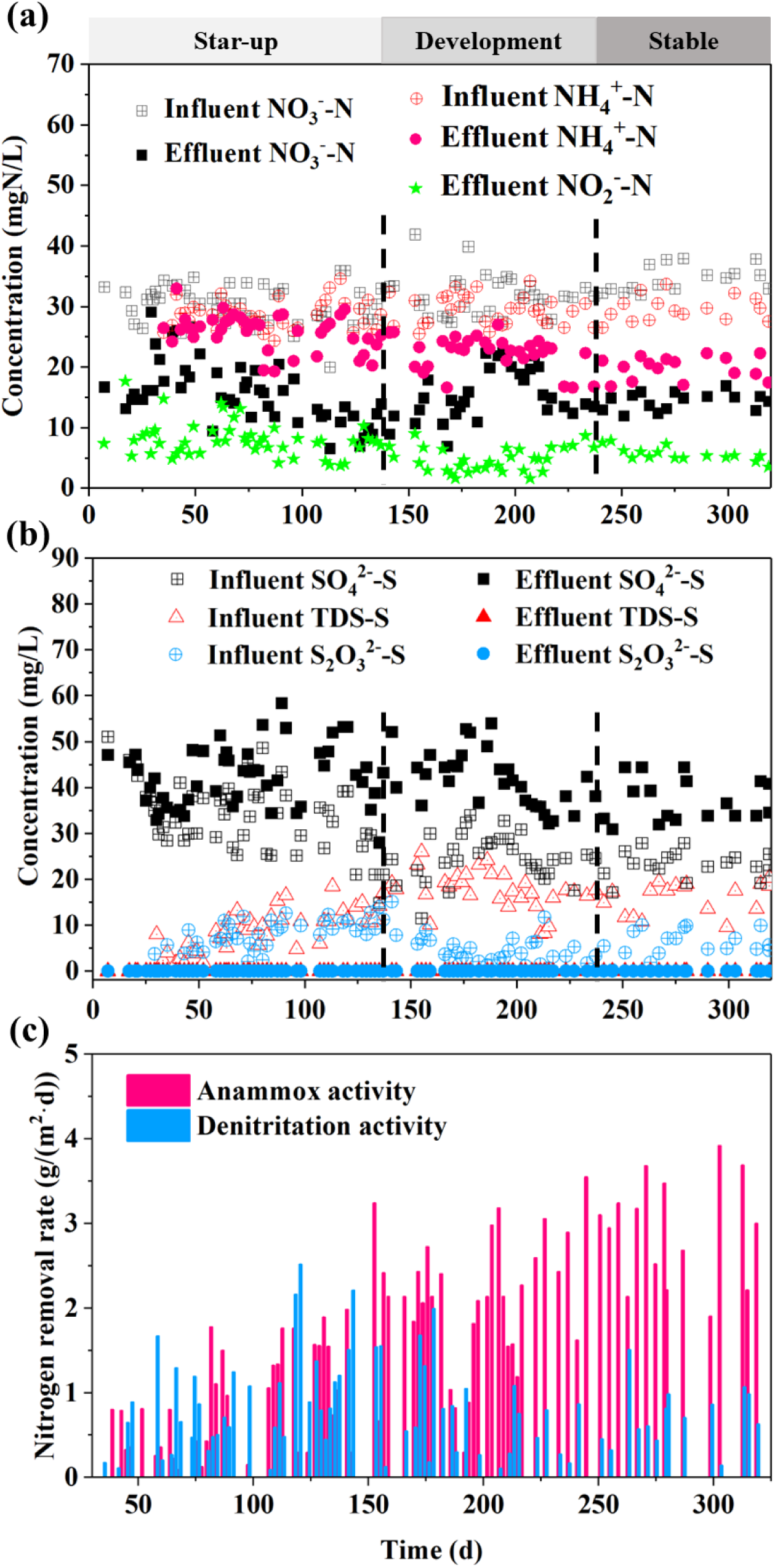
MSPDA-MBBR performance of nitrogen and sulfur transformation in Period I: conversion of a) nitrogen compounds, b) sulfur compounds, c) contribution of anammox and denitrification to nitrogen removal in MSPDA-MBBR (Note: magnetic stirrer has a stirring problem started on day 190, then replaced by a new one on day 199).

#### 3.1.2 Evaluation of TIN removal via Denitrification and anammox

Based on the routine data, the contribution of denitrification and anammox to TIN removal in MSPDA-MBBR were analyzed. Meanwhile, the activities of denitritation and anammox (including removal of ammonium and nitrite) were discussed.

During days 98−135, 9.1 mgN/L and 7.3 mgN/L of TIN removal were determined via anammox and denitrification, respectively based on the performance data. The result suggests the establishment of anammox and autotrophic denitrification. It is worth noting that the anammox biomass was normally inoculated in mainstream anammox process directly or indirectly (simultaneous provision of ammonium and nitrite) previously (Deng et al., 2021a; Du et al., 2017; Kalyuzhnyi et al., 2006). With neither of the strategy applied in this MSPDA-MBBR, anammox activity was observed within 135 days, comparable to the typical range of 60−130 days for start-up of anammox process (Deng et al., 2019; Tang et al., 2010; Wang et al., 2009b). After achievement of stable performance during days 271–319, anammox and denitrification contribute to TIN removal of 19.5 mgN/L and 4.7 mgN/L, respectively. Meanwhile, with residual nitrite of 5.8 ± 1.1 mgN/L and the sulfur loss of 28%, the total TIN removal could be further improved by fully utilization of produced nitrite and electrons contained in sulfur compounds. The potential of TIN removal was 39.3 mgN/L attributed to that (1) all nitrite removed via anammox would contributed to extra 10.3 mgN/L TIN removal; (2) noted that the 28% of sulfur loss (calculated as S^0^) containing electrons constitutes to 20% of total utilized electrons, full utilization of the lost electrons would contribute to another TIN removal of 3.9 mgN/L and 0.9 mgN/L through anammox and denitrification, respectively.

Meanwhile, the surface-specific denitritation rate decreased from 1.3 gN/(m^2^·d) to 0.7 gN/(m^2^·d) during days 136–319 as shown in Fig. 2c. The anammox rate continuously grew from 1.4 gN/(m^2^·d) into 2.8 gN/(m^2^·d). Therefore, the ratio of nitrogen removal via anammox to TIN removal achieved 81% in this study, which is comparable to 80−90% in SPDA process with external dosage of thiosulfate or sulfide (Deng et al., 2019; Deng et al., 2021b). Notably, the volumetric anammox rate achieved as high as 0.7 gN/(L·d), close to the maximum rate of 0.3−0.6 gN/(L·d) obtained in mainstream wastewater treatment (Laureni et al., 2016). These results demonstrate that high anammox activity could be developed in MSPDA without external dosage of sulfide or thiosulfate, even when operated at a low HRT of 0.7 h and room temperature of 25℃.

#### 3.1.3 Enrichment and dynamics of functional bacteria in MSPDA-MBBR

The relative abundances of dominant bacteria in MSPDA-MBBR were analyzed and results are presented in Fig. 4. At phylum level, *Proteobacteria* dominants with abundance of 63.7%, following by *Bacteroidota, Actinobacteriota, Desulfobacterota, Chloroflexi, Firmicutes, Campilobacterota* and *Planctomycetes* with abundances of 9.2%, 5.4%, 5.3%, 4.9%, 2.9%, 2.3% and 1.7% respectively after 106 days’ cultivation. Among which, SOB as the functional bacteria for autotrophic denitrification, were mainly found belong to *Proteobacteria* (Kostrytsia et al., 2018), and recently uncultured *Desulfobacterota* role in denitrification was also revealed (Langwig et al., 2022). The abundances of *Proteobacteria* and *Desulfobacterota* decreased to 38.0% and 2.4% on day 319 (Fig. 4b). Meanwhile, *Planctomycetes* increased from 0% to 4.3% during 319 days’ cultivation. In genus level, total abundance of denitrifying bacteria, including autotrophic denitrifiers (SOB) *Thiobacillus (*β- proteobacteria) and *Sulfurimonas (*γ-Proteobacteria), heterotrophic denitrifiers like *Denitratisoma* and *Flavobacterium*, and facultative denitrifiers *Thauera,* was 17.1% on day 106 which continuously decreased to 7.1% on day 240 and then to 3.8% on day 319. *Thiobacillus, Sulfurimonas,* and *Thauera* are frequently found in SPDA and PD/A process (Deng et al., 2021b; Ma et al., 2017). Besides, *Thiobacillus* and *Sulfurimonas* could utilize sulfide and thiosulfate for fast denitratation (Qian et al., 2021). Meanwhile, enrichment of *Thauera* which adapts to both organics and sulfide could also facilitate fast denitratation (Liang et al., 2020). The enrichment of these bacteria is in line with the fast nitrite provision obtained in this study. Meanwhile, anaerobic ammonium oxidizing bacteria (AnAOB) were successfully enriched on day 106 as presented in Fig. 4c. On day 106, the abundances of *Candidatus Scalindua* and *Candidatus Kuenenia* were 0.6% and 0.2% which then increased to 1.4% and 1.8%, respectively on day 319. The *Candidatus Kuenenia* as a K-strategist with high affinity for substrate (Kosgey et al., 2020) was dominant in this system, supporting the development of anammox under low nitrite concentration. Meanwhile, *Candidatus Scalindua* commonly found in marine environment (Awata et al., 2013) was also enriched in this study, possible due to 5% seawater addition of the influent for the supply of sulfate. In summary, the decrease in denitrifiers and increase in AnAOB during 319 days, were corresponding to increased ratio of nitrogen removal via anammox to total nitrogen removal as describe in section 3.1.2. Meanwhile, it indicates the occurrence of resource partitioning in which nitrate was utilized by denitrifying bacteria and produced nitrite was used by AnAOB, beneficial to the long coexistence of SOB and AnAOB.

**Fig. 4.**
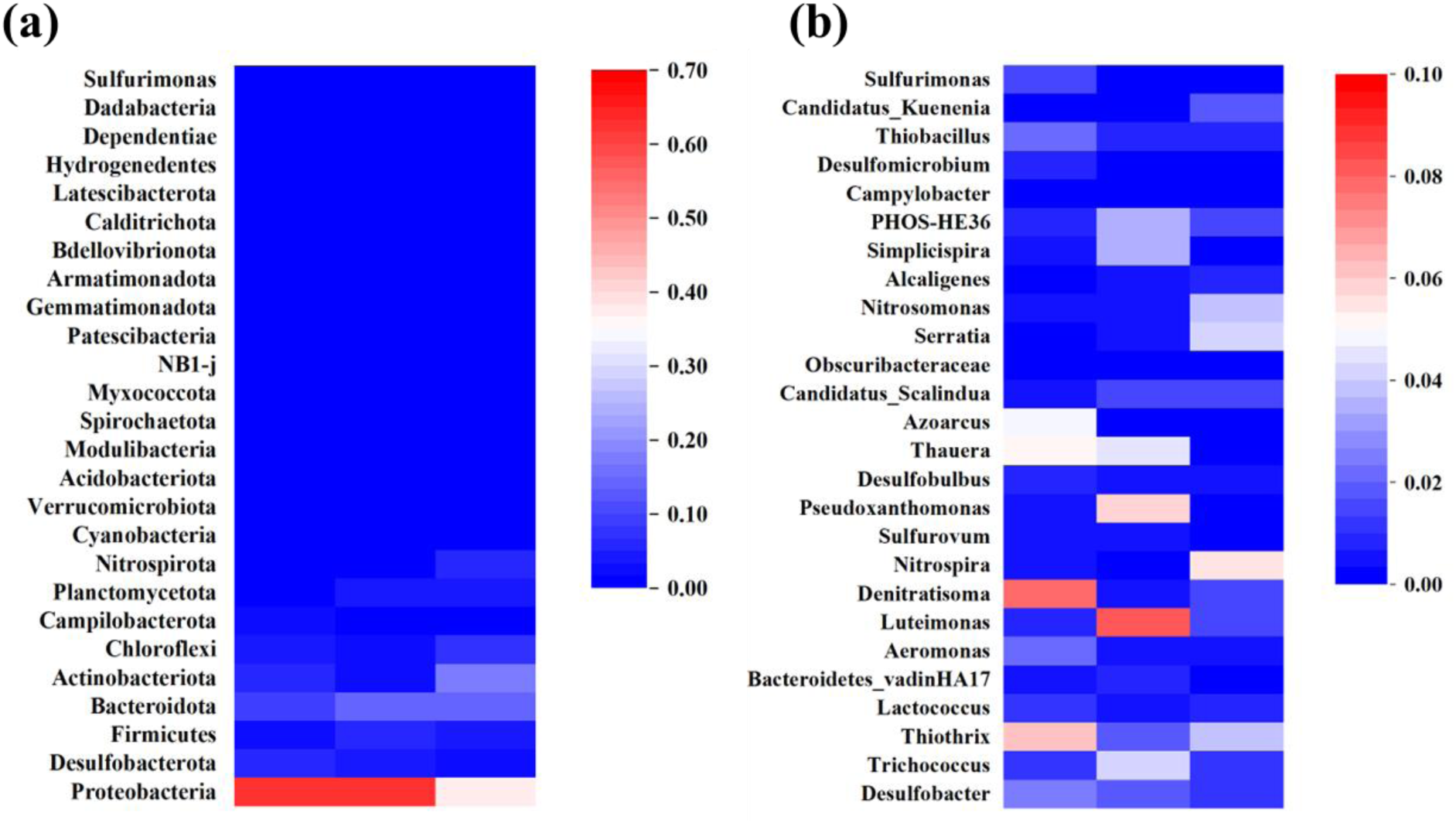
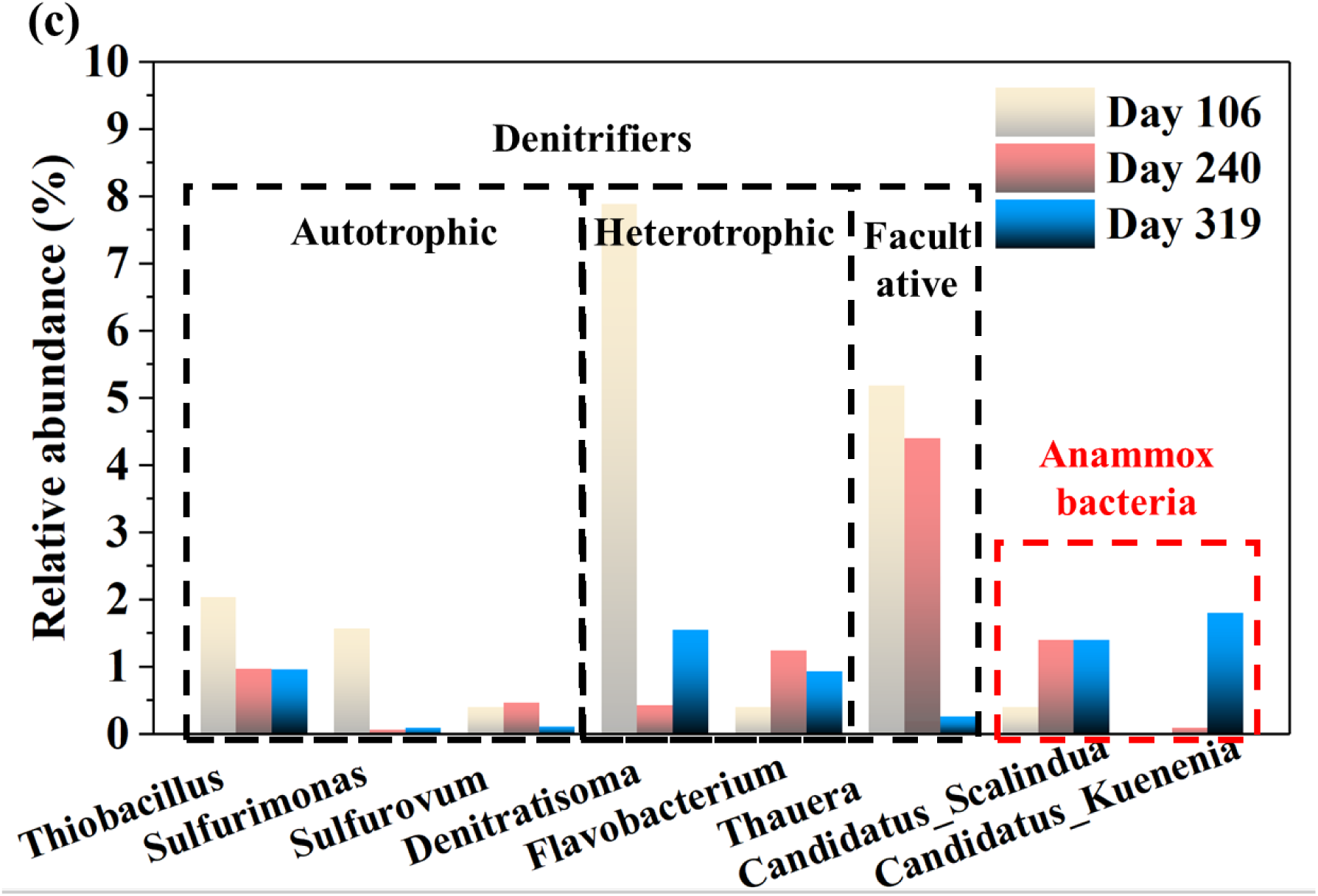
Microbial community dynamics in MSPDA-MBBR on day 106, day 240, and day 319, respectively: (a) phylum level; (b) genus level; (c) abundance of functional bacteria.

### 3.2 S and N conversions in MSPDA

With the high-rate MSPDA developed in this study, the conversion behaviours of S and N, which might relate to nitrite accumulation rate hence the anammox rate, is yet to be investigated. Although SPDA has been demonstrated with using sulfide or thiosulfate as the single electron donor (Deng et al., 2022), the S and N conversions and interactions in MSPDA with coexistence of sulfide and thiosulfate are to be elucidated. Therefore, batch A−D were conducted for further investigation.

#### 3.2.1 TdS-driven denitrification

Batch A was performed to determine sulfide-driven denitrification activity, and the results are presented in Fig 5a. TdS was quickly oxidized to elemental sulfur which was then slowly oxidized to sulfate as reported (Xu et al., 2016). Therefore, the batch test was divided into stage 1 (0−30 min) and stage 2 (30−360 min). In stage 1 with initial TdS concentration of 27.6 ± 0.1 mgS/L, TdS was rapidly decreased to 0.3 ± 0.1 mgS/L in 30 min with a surface-specific oxidation rate of 546.1 mgS/(m^2^·h). Sulfate was not detected during the first 30 min, and it slowly increased to 22.1 ± 0.2 mgS/L at the end of stage 2 with a rate of 33.6 mgS/(m^2^·h). The rates of TdS oxidation and sulfate generation were close to the typical range of 188−484 mgS/(m^2^·h) and 4−49 mgS/(m^2^·h), respectively (Cui et al., 2019c). After 360 mins’ reaction, 103% of sulfur mass balance was achieved. Correspondingly, 31.6 ± 0.8 mgN/L of nitrate rapidly decreased with a denitratation rate of 226.8 mgN/(m^2^·h) in stage 1, much higher than 35.1 mgN/(m^2^·h) of denitritation rate. The results indicate that a fast nitrite accumulation rate of 191.7 mgN/(m^2^·h) was achieved with a high nitrite accumulation ratio of 85%. Meanwhile, the nitrite accumulation rate in this stage is 6.2−45.7 times of it reported in previous sulfide-oxidizing autotrophic denitrification (Cui et al., 2019c). The high nitrite accumulation rate revealed in stage 1 supports the high anammox activity developed in MSPDA-MBBR. In stage 2 with S^0^ slowly converted to sulfate, and denitratation and denitritation rate decreased to 26.9 mgN/(m^2^·h) and 12.9 mgS/(m^2^·h) with nitrite accumulation ratio of 52%. The nitrite accumulation rate of 14.0 mgN/(m^2^·h) here, much slower than it in stage 1, was in the reported range of 4.2−31.8 mgN/(m^2^·h) (Cui et al., 2019c; Koenig and Liu, 1996). Moreover, the electrons flew from sulfide/S^0^ to nitrate/nitrite were analyzed in both stage 1 and stage 2 based on the reduction of nitrate and nitrite. It was found that denitratation utilized 81% and 58% of the electrons in stage 1 and stage 2, respectively. During this whole batch, the electron balance was 107% based on the initial and final concentrations of S and N compounds.

**Fig. 5.**
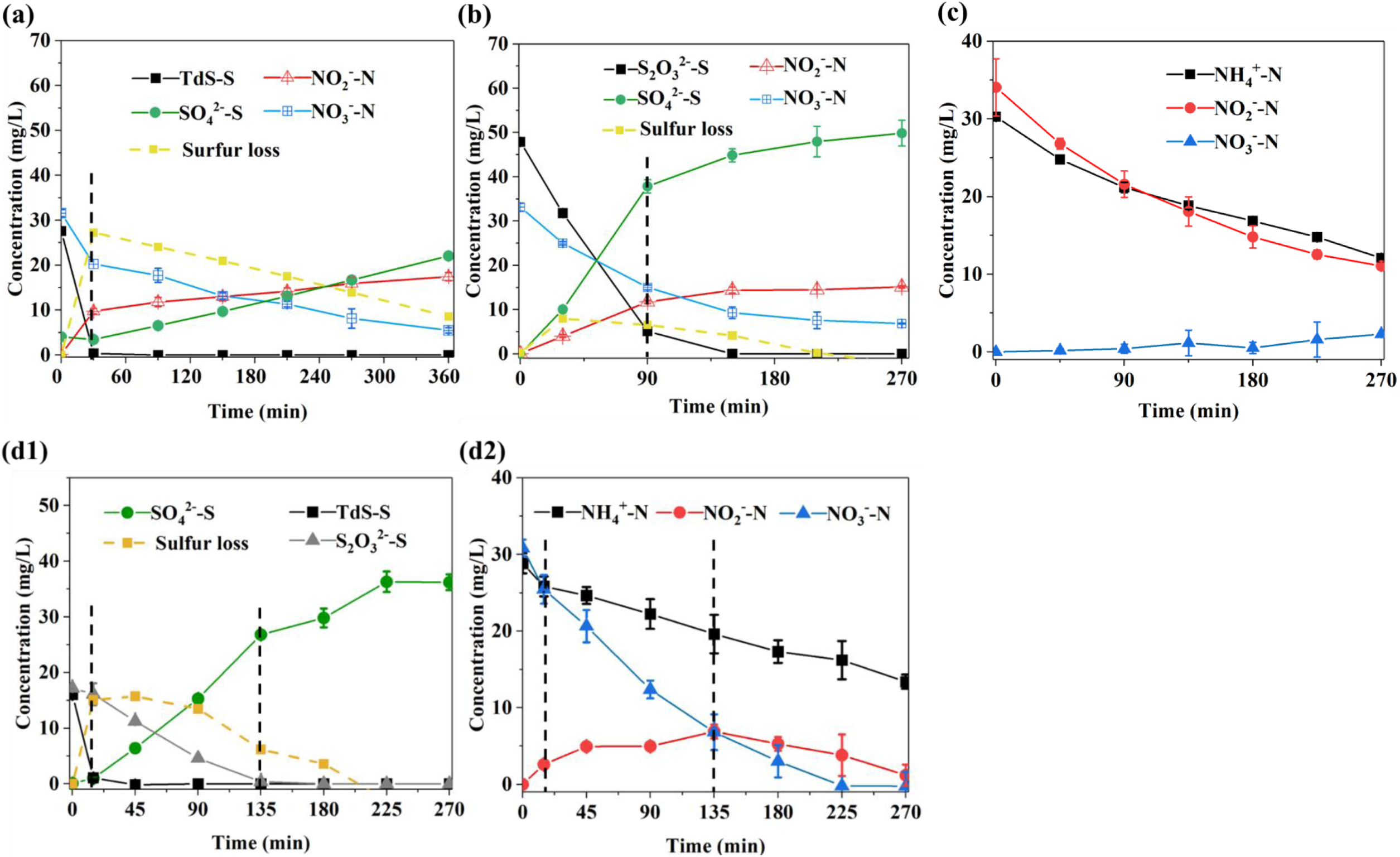
Investigation of MSPDA in batch tests: a) Batch A, denitrification with TdS as sole electron donor; b) batch B, denitrification with thiosulfate as the only electron donor; c) batch C, anammox with ammonium and nitrite provision; d1) and d2) S and N profiles for batch D, MSPDA with TdS and thiosulfate as mixed electron donors.

**Fig. 6.**
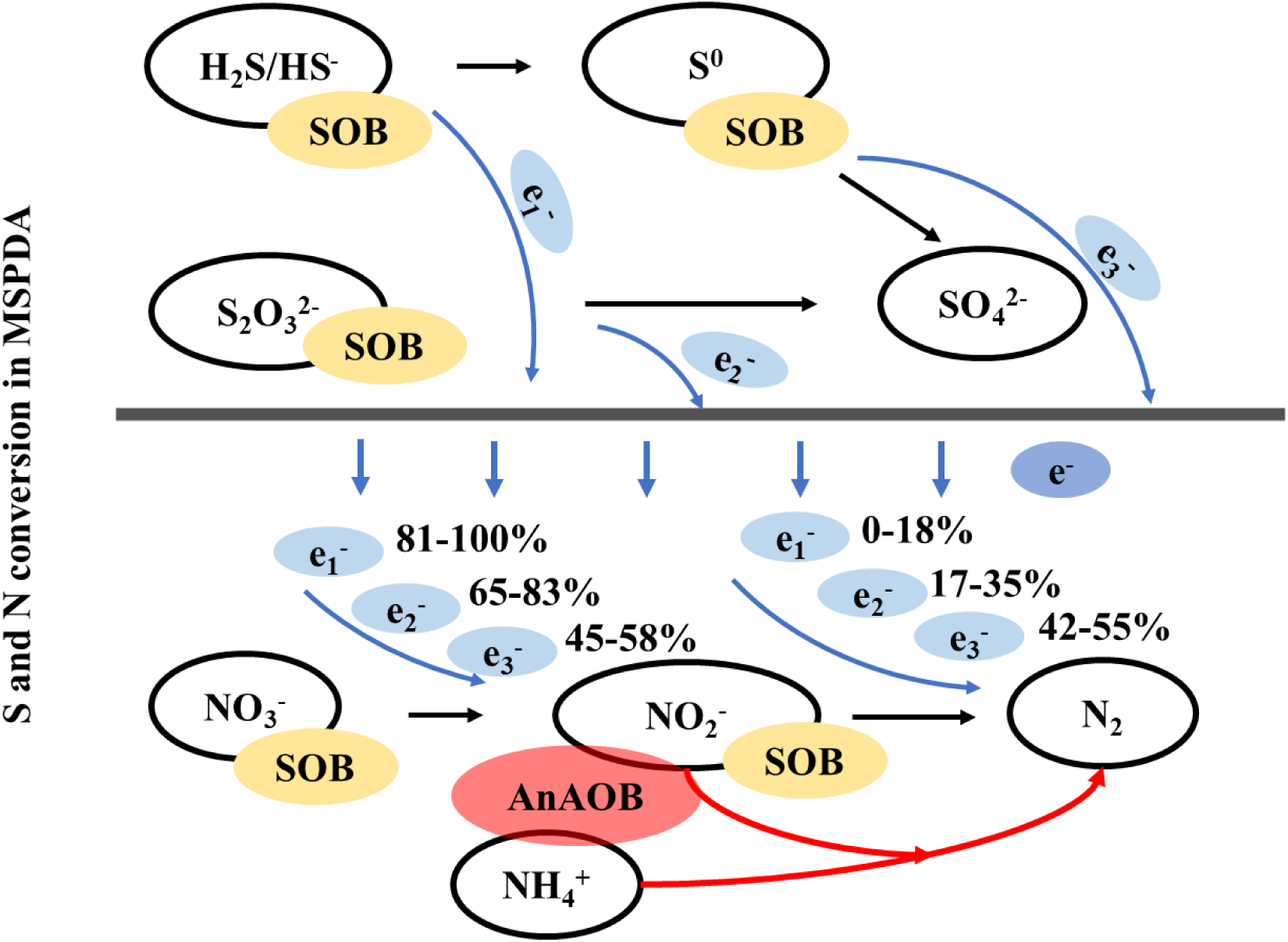
Proposed mechanism of sulfur and nitrogen compounds conversion in MSPDA (e_1_^-^, e_2_^-^, and e_3_^-^, refer to electrons obtained from oxidation of sulfide, thiosulfate, and S^0^ respectively).

#### 3.2.2 Thiosulfate-driven denitrification

Batch B was performed to determine thiosulfate-driven denitrification activity in MSPDA. Similar to batch A, the test was separated into stage 1 (0−90 min) and stage 2 (90−210 min) as presented in Fig 5b. In stage 1, 47.9 ± 0.1 mgS/L thiosulfate was decreased fast with an oxidation rate of 285.1 mgS/(m^2^·h). Meanwhile, sulfate produced with a rate of 252.5 mgS/(m^2^·h). Different from half thiosulfate converted to S^0^ in a thiosulfate-driven PDA bioreactor (Deng et al., 2019), thiosulfate in this study were mainly oxidized to sulfate without intermediate formation. In stage 2, with the left 5.1 ± 0.7 mgS/L of thiosulfate disappeared at 150 min, sulfate concentration increased to 44.9 ± 1.5 mgS/L with 94% of sulfur balance. For nitrogen conversion with initial nitrate concentration as 33.1 ± 0.9 mgN/L, denitratation rate and denitritation rate in stage 1 were 120.6 mgN/(m^2^·h) and 42.6 mgN/(m^2^·h), respectively. Therefore, the nitrite accumulation rate was 78.0 mgN/(m^2^·h). Then in stage 2, 5.7 mgN/L decrease of nitrate and 2.4 mgN/L increase of nitrite were obtained at 150 min. Without thiosulfate during 150−270 min, further 2.4 mgN/L of decrease in nitrate and 0.7 mgN/L of increase in nitrite might be due to existence of S^0^ derived from original carriers in bioreactor (Cui et al., 2019c). The nitrite accumulation ratio in stage 1 was 65% with thiosulfate as an electron donor. Denitratation consumed 65% of total electrons donated by thiosulfate oxidation (S_2_O_3_ ^2-^ ➔ SO_4_^2-^). Moreover, compared to batch A, 89% of electrons in thiosulfate were donated in only 90 mins, leading to that utilization rate of electrons in thiosulfate was 3.6 times higher than sulfide. As electrons flows fast from thiosulfate to nitrate, thiosulfate existence may contribute to the fast denitrification. The balances of sulfur mass and electron were achieved 104% and 102% respectively after 210 mins’ reaction.

#### 3.2.3 Anammox

Batch C was performed to further determine the anammox activity. Ammonium and nitrite were decreased to 12.1 ± 0.6 mgN/L and 11.1 ± 0.3 mgN/L with initial concentrations of 30.3 ± 0.2 mgN/L and 34.0 ± 3.7 mgN/L as Fig. 5c shows. In batch C, 2.3 ± 0.1 mgN/L nitrate was produced within 270 mins. Spontaneous removal of ammonium and nitrite demonstrated the anammox reaction mediated by the living carriers taken from MSPDA-MBBR. Meanwhile, the ratio of nitrite consumption/ammonium consumption is 1.27 close to reported value 1.32 for anammox reaction (Strous et al., 1998). The conversion rate of ammonium and nitrite were 40.5 mgN/(m^2^·h) and 50.8 mgN/(m^2^·h) with nitrate production rate of 5.8 mgN/(m^2^·h). The nitrite utilization rate by anammox was much lower than the nitrite accumulation rates of 78.0−191.7 mgN/(m^2^·h) obtained in stage 1 of batch A and B, indicating the sufficient provision of nitrite for anammox development. Meanwhile, compared to denitritation rates of 12.9−42.6 mgN/(m^2^·h) obtained in batch A and B, the nitrite utilization rate in anammox was the highest, indicating that anammox could compete over denitritation for nitrite. R^2^ for all specific rates in batch A, B, and C are higher than 0.9.

#### 3.2.4 MSPDA

Batch D was designed to further reveal S and N conversion in MSPDA process. According to results from batch A, B and D, oxidation of sulfide and thiosulfate in batch D was divided into three stages: stage1 (0−15 mins), sulfide was oxidized at first with S^0^ formation; stage 2 (15−135 mins), formed S^0^ and thiosulfate was oxidized to sulfate, during which thiosulfate was consumed faster than S^0^; stage 3 (135−270 mins), the residual S^0^ was oxidized to sulfate. In stage 1, initial concentrations of TdS and thiosulfate were 16.0 ± 0.1 mgS/L and 17.2 ± 0.3 mgS/L, respectively. Specific oxidation rate of TdS as shown in Table 3, was 597.5 mgS/(m^2^·h). After sulfide depletion, thiosulfate oxidation rate increased to 84.0 mgS/(m^2^·h) with sulfate production rate as 129.0 mgS/(m^2^·h) during stage 2. At last, in stage 3, thiosulfate was completely consumed, and sulfate generation rate decreased to 63.3 mgS/(m^2^·h) with sulfur balance of 109%. The denitratation rates of 216.0 and 83.0 mgN/(m^2^·h) in stage 1 and stage 2 were close to the rates obtained in batch A and B during oxidation of sulfide and thiosulfate, respectively. Ammonium removal rate via anammox was 120.0 mgN/(m^2^·h) in stage 1, much higher than the rate obtained in batch C. Therefore, the low concentration of sulfide indeed promoted annamox reaction rate as reported (Van de Graaf et al., 1996). This could be due to that the fast oxidation of sulfide could cleavage the DO fast with generation of strict anaerobic conditions thus enhancing denitrification and anammox (Wenk et al., 2013). After stage 1, ammonium removal rate decreased to 29.0 mgN/(m^2^·h) during stage 2, and maintained at 27.5 mgN/(m^2^·h) finally in stage 3. Notably, accumulation of nitrite was found in both stage 1 (2.6 mgN/L) and stage 2 (4.3 mgN/L) with the oxidation of sulfide and thiosulfate, respectively. And the accumulation of nitrite supported the anammox reaction rate maintained at 27.5 mgN/(m^2^·h) in the following stage 3 with nitrite sharply decreased to 1.2 ± 1.2 mgN/L, indicating that nitrite production in oxidation of sulfide (S^2-^ ➔ S^0^) and thiosulfate was critical for the high anammox activity. In total, the utilization rate of electron donor in MSPDA with 35% of electron donor provided by thiosulfate was still promoted by 43% compared to it in TdS-driven denitrification. Meanwhile, the electron usages for denitration and denitritation were determined based on the stoichiometric ratios obtained in batch C. 100%, 83%, and 45% of electrons in sulfide, thiosulfate, and S^0^ were found utilized in denitratation with an overall electron balance of 110% in this batch.

**Table 3.**
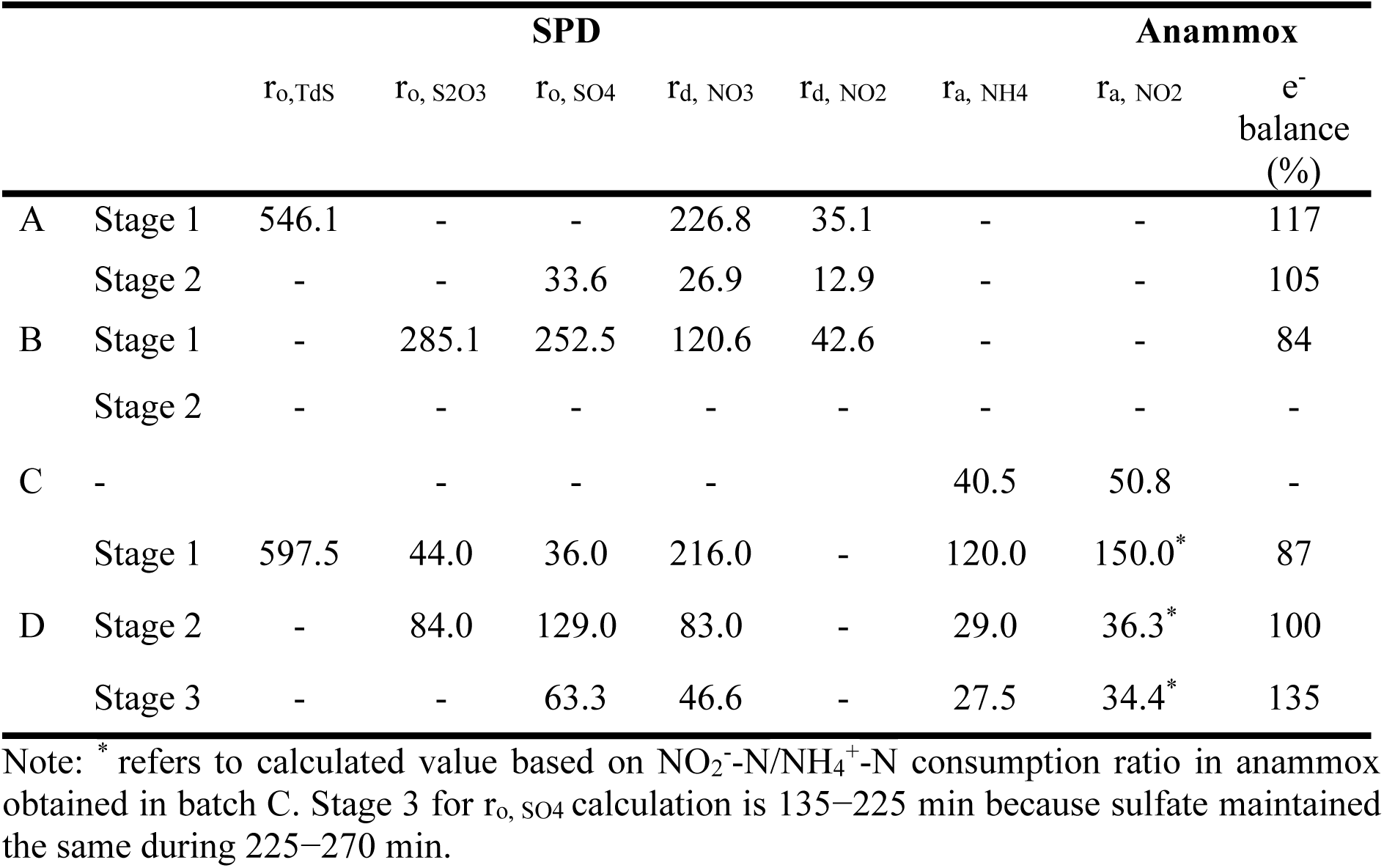
Kinetic rates obtained from batch tests A-D (unit: mgN/(m^2^·h) or mgS/(m^2^·h))

#### 3.2.5 Speculation of S and N conversion

To further analyze S and N conversion in MSPDA, the conversion mechanism was speculated, based on batch A−D results and related thermodynamics.

The S and N conversion was depicted in Fig. 4 based on results from batch A, B and D. Sulfide was oxidized fast and SO_4_^2-^ was only produced after sulfide disappearance as results shown in both batch A and batch D, indicating sulfide was oxidized (S^2-^ ➔ S^0^ ➔ SO_4_^2-^) as described in previous sulfide-driven denitrification and SPDA processes (Deng et al., 2021b; Shao et al., 2010). Corresponding to this sulfide oxidation pathway, both β- and γ-proteobacteria enriched in this bioreactor contain sulfide-quinone reductase which oxidizes sulfide with S^0^ formation (Ghosh and Dam, 2009). Notably, nitrite accumulation during TdS oxidation to S^0^ here is in line with nitrite production with S/N ratios of 0.5−5.6 using an oil reservoir microbial culture (An et al., 2010), while different from no nitrite production with S/N ratio between 0.7−3.8 applying the biofilm enriched in a sulfide-driven denitrification bioreactor (Cui et al., 2019c). The difference is possibly due to the upregulation of genes related to denitrification including nar, nir, and nor genes which are associated with nitrate reductase, nitrite reductase, and nitric oxide reductase, respectively under different conditions (Beller et al., 2006). After sulfide depletion, thiosulfate as the main electron donor was converted to sulfate directly (S_2_O_3_^2-^ ➔ SO_4_ ^2-^) as indicated in batch B and D. This is in accordance with that *Thiobacillus* and *Sulfurimonas* enriched in this study were capable of oxidizing thiosulfate to sulfate directly (Shao et al., 2010). This pathway has already been proposed as alphaproteobacterial Sox pathway governed by the Sox complex. In which, the sulfane sulfur of thiosulfate was oxidized via complex reactions while the sulfone sulfur is directly oxidized and released as sulfate (Quentmeier et al., 2003).

Notably in MSPDA, 81−100%, 65−83%, and 45−58% of electrons from sulfide (S^2-^➔ S^0^), thiosulfate, and S^0^, respectively were utilized for denitratation based on all batch tests results. Meanwhile, the sulfur-driven denitratation rates tend to decrease with the different electron donors in the order of TdS > thiosulfate > S^0^ in this study. This could be supported by Eqs. (1), (2) and (4), with the sequence of Gibs free energy release among oxidation of sulfide, thiosulfate and S^0^ from the highest to the lowest. Moreover, 59% of the nitrite in total was removed via anammox, suggesting that anammox could compete with autotrophic denitritation in MSPDA. This is in accordance with that anammox reaction (Kartal et al., 2011) could release more or comparable Gibs energy compared to S^0^- or thiosulfate-based denitrification as presented in Eqs. (2)−(6).

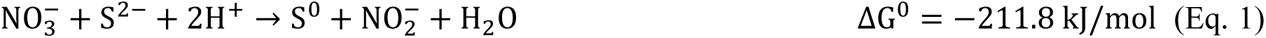

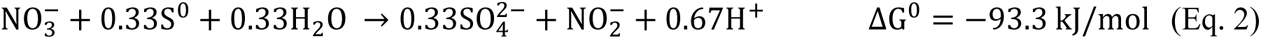

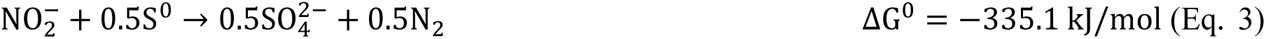

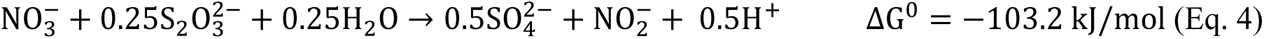

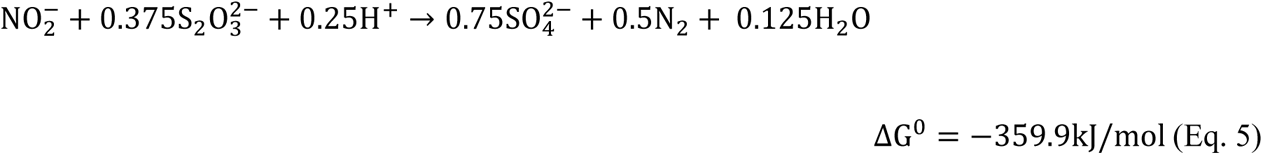

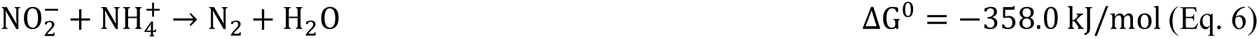

### 3.3 SANIA in Period II: performance of the SANIA system

Although MSPDA system was successfully established, integration of the real nitrifying effluent into the SANIA process is necessary for two uncertainties caused by residual DO in nitrifying effluent: (1) it would consume the valuable and limited electron donors (TdS and thiosulfate), causing deficiency of electron donors for denitratation; (2) it might increase the DO in MSPDA-MBBR thus affecting the activity of anammox (Strous et al., 1997).

To further verify the feasibility of SANIA system, nitrification MBBR (N-MBBR) with a DO of 2.7 ± 0.9 mg/L was applied for nitrite or nitrate provision during days 325−455 (Period II). Performances of ERATO-, MSPDA, N-MBBRs in SANIA were presented as Fig. 7 and Table 4. After day 387, performance of three bioreactors become stable. In ERATO-MBBR, fed influent with DO of 2.7 ± 0.9 mg/L, 15.4 ± 1.3 mg/L of TdS and 6.1 ± 2.5 mg/L of thiosulfate were produced. Surface-specific sulfate reduction rate and COD removal rate stabilized at 0.4 gS/(m^2^·d) and 1.0 g/(m^2^·d), respectively. In MSPDA-MBBR with HRT prolonged to 1.0 h, sulfur balance achieved 92% which was higher than 72% obtained in Period I, indicating the full utilization of sulfur compounds. With nitrified effluent recycled for nitrate supply, the ORP in MSPDA-MBBR increased from −234 mV to 50 mV, indicating a higher DO obtained in this circumstance. Still, the ammonium and nitrate were effectively removed with TIN decreased from 33.0 ± 2.1 mgN/L in influent to 13.9 ± 1.8 mgN/L in this bioreactor. Specifically, nitrate was reduced in MSPDA-MBBR with only 1.3 ± 0.9 mgN/L in effluent which means the electron supply is sufficient for complete nitrate reduction. According to ammonium removal in MSPDA-MBBR, anammox contributed to 74% of removed TIN. With increased ORP in MSPDA-MBBR, anammox reaction rate of 0.4 gN/(L·d) still achieved in the SANIA process within the typical range of 0.3−0.6 gN/(L·d) (Laureni et al., 2016). In contrast, 10.3 ± 1.3 mgN/L of ammonium and 2.3 ± 0.9 mgN/L of nitrite in effluent indicated the further improvement of nitrogen removal via anammox could be achieved via prolonging the reaction time. In short, even with the residual DO of 2.7 ± 0.9 mg/L in nitrifying effluent and a recycle ratio of 2.0, the obtained nitrogen removal performance and the practical reaction rate in MSPDA demonstrated its applicability for mainstream wastewater treatment. In N-MBBR, which is the last step of SANIA process, ammonium was effectively oxidized with ammonium, nitrite, and nitrate in influent maintained at 2.1 ± 0.9 mgN/L, 2.0±1.0 mgN/L, and 8.7 ± 1.8 mgN/L respectively. Nitrate concentration obtained here was quite close to the theoretical result in stable operation as calculated in SI 2.2. Therefore for the whole system, COD, ammonium, and TIN removal efficiency were reached 90%, 93% and 61% separately with COD, ammonium, and TIN below 10 mg/L, 2.1 ± 0.9 mgN/L, and 13 mgN/L, respectively. In short, the feasibility of the integrated SANIA system was demonstrated via over 130 days’ operation in this study.

**Fig. 7.**
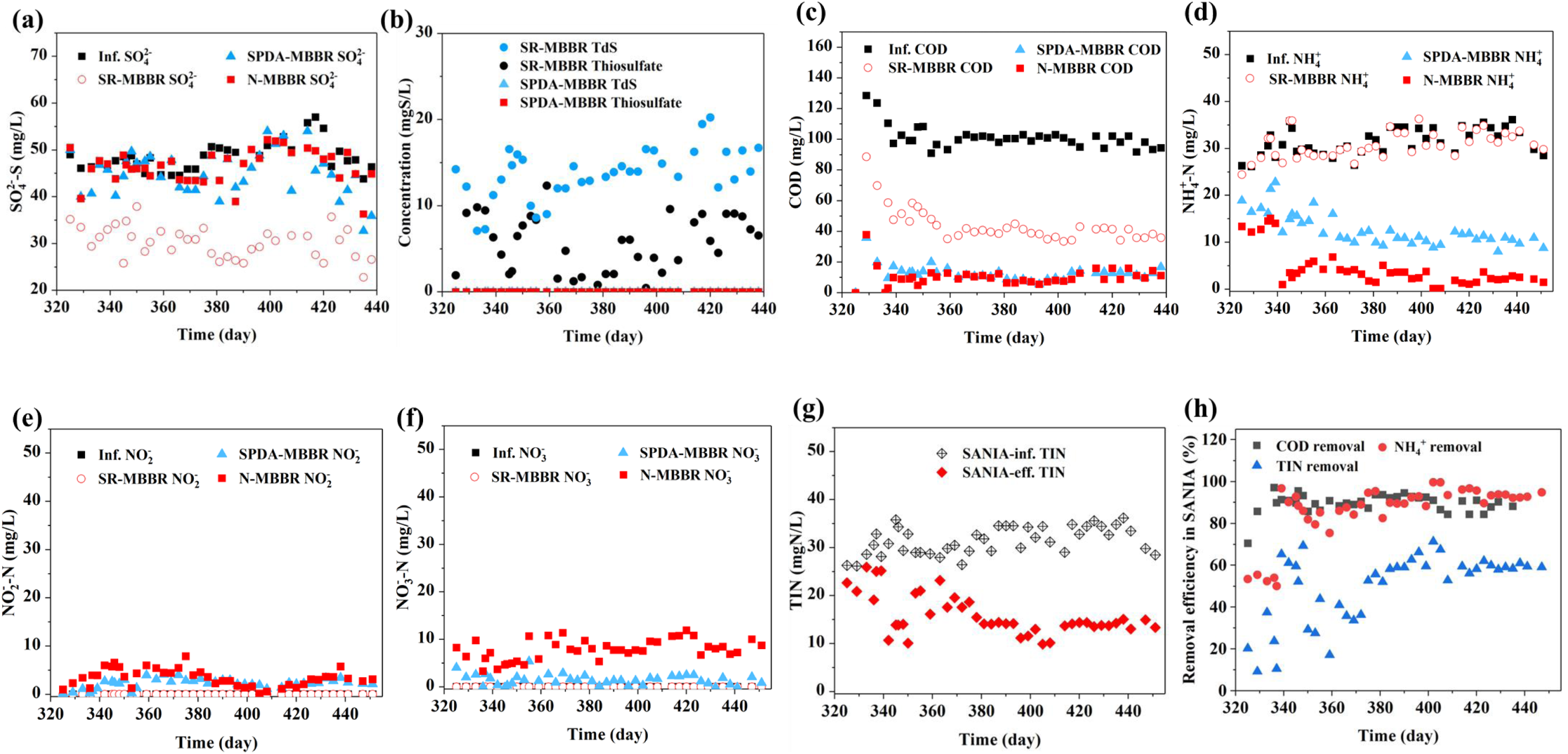
Performance of SANIA process: profile of a) SO_4_^2-^; b) TdS and thiosulfate; c) COD; d) NH_4_^+^; e) NO_2_^-^; f) NO_3_^-^; g) TIN; h) removal efficiency of COD, NH_4_^+^ and TIN.

**Table 4.** Performance for SANIA system in Period II.

| Parameters | Influent | ERATO-MBBR | MSPDA-MBBR | N-MBBR |
| --- | --- | --- | --- | --- |
| Sulfate (mg S/L) | 50.7±3.1 | 29.6±2.9 | 46.5±4.7 | 48.3±3.3 |
| Sulfide (mg S/L) | - | 15.4±2.3 | 0.0 | 0.0 |
| Thiosulfate (mg S/L) | - | 6.1±2.5 | 0.0 | 0.0 |
| Organics (mg COD/L) | 99.5 ± 3.4 | 38.7±3.7 | 10.7±2.7 | 9.9±3.5 |
| Ammonium (mg N/L) | 33.0±2.1 | 32.3±2.4 | 10.3±1.3 | 2.1±0.9 |
| Nitrite (mg N/L) | - | - | 2.3±0.6 | 2.0±1.0 |
| Nitrate (mg N/L) | - | - | 1.3±0.9 | 8.7±1.8 |
| Alkalinity (mg/L) | 205 ±12 | 273±31 | 153±16 | 99±17 |
| TSS in effluent (mg/L) | - | - | - | 20.2±4.4 |
| VSS in effluent (mg/L) | - | - | - | 16.7±4.2 |
| TIN (mg/L) | 33.0±2.1 | 32.3±2.4 | 13.9±1.8 | 12.9±1.7 |
| Sulfur balance (%) | - | 100 | 92 | 95 |
| TIN removal via anammox (%) | - | - | 74 | - |

Note: the data obtained during days 387-455 were used for the calculation.

### 3.4 Evaluation of SANIA process

Compared to conventional activated sludge process, in which ammonium is all oxidized to nitrate with an typical HRT of 10−24 h (Ekama and Wentzel, 2020), the SANIA process developed in this study could save 50−81% space and 20% of aeration energy for nitrogen removal. Meanwhile with only 16.7 ± 4.2 mg/L VSS production in effluent, sludge yield is only 0.19 gVSS/gCOD with due to low sludge yield of SRB, SOB, AnAOB, and nitrifying bacteria. The effluent SS concentration of 18 mg/L is much lower than 150−250 mg/L obtained in typical MBBR process for treating mainstream wastewater (Kängsepp et al., 2020). Moreover, the alkalinity consumption was only 106 mg/L with effluent pH of 7.2 ± 0.1, indicating no demand of external alkalinity. Furthermore, by combining this process with organics recovery, energy-positive treatment of wastewater is possible. For instance, when CEPT is applied for organics harvest as shown in Fig. 8, the implementation of SANIA in the main line would need an energy input of 0.23 kWh/m^3^ wastewater (based on parameter values in SI 7). This low energy consumption is mainly caused by that 60% of aeration energy input for organics capture/removal and nitrogen removal is saved. Meanwhile, captured COD in form of sludge contains higher energy than secondary sludge (Wan et al., 2016). Through anaerobic digestion, cogeneration, and incineration for energy recovery (Abuşoğlu et al., 2017; McCarty et al., 2011; Wan et al., 2016), the captured COD could produce 0.30 kWh/m^3^ of energy. Therefore, the energy yield is 0.07 kWh/m^3^ wastewater, indicating the achievement of energy-neutral wastewater treatment by application of SANIA.

**Fig. 8.**
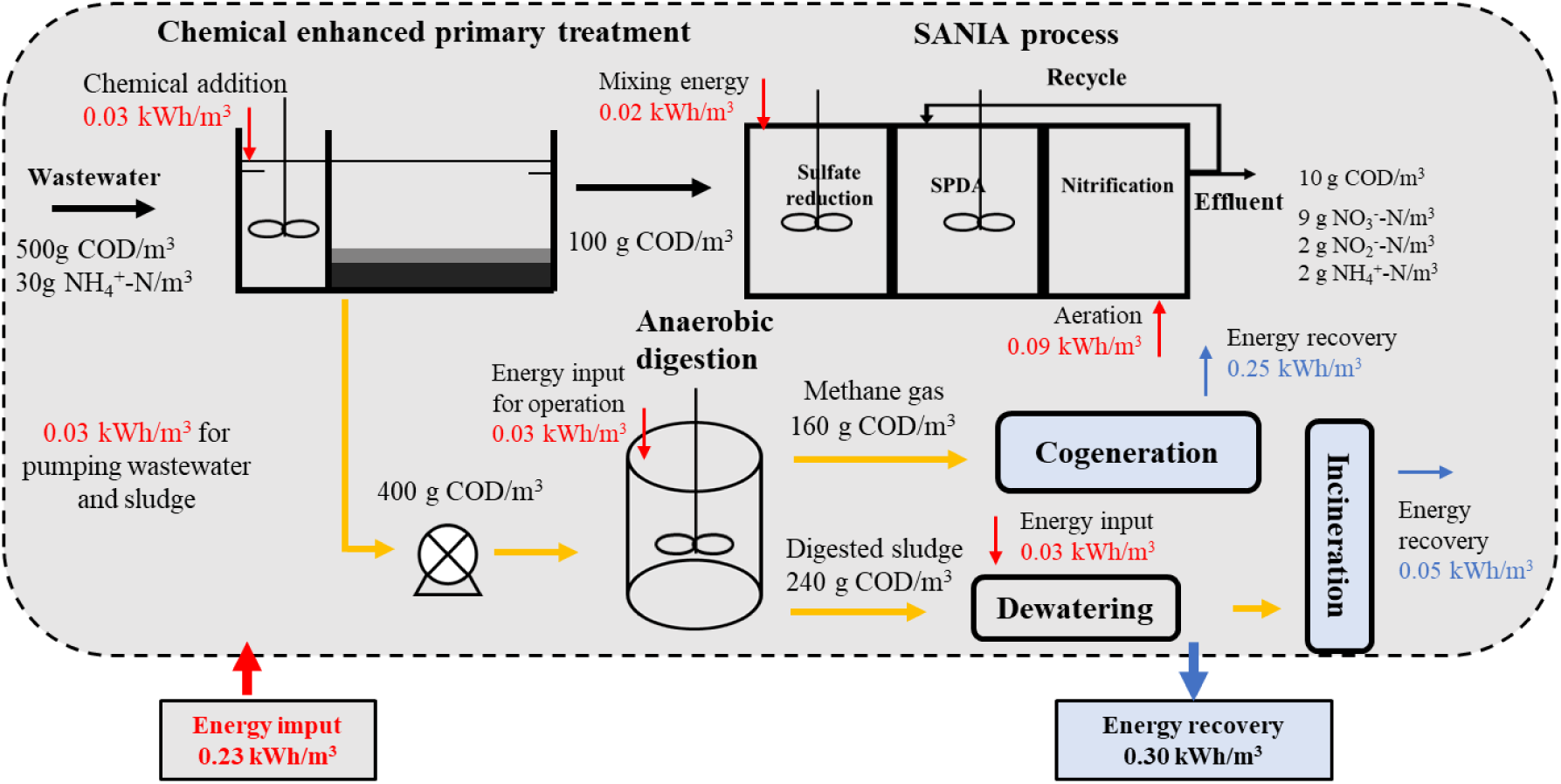
Energy analysis based on the performance of SANIA process combining chemical enhanced primary treatment, the calculation was based on values summarized in Table S5.

## 4. Conclusion

The novel SANIA process was successfully developed in this study, providing a promising option for anammox application in mainstream wastewater treatment. Main conclusions are summarized as below:

- Without anammox biomass inoculation, MSPDA was successfully developed with a high surface-specific anammox rate of 2.8 gN/(m^2^·d) and 81% of the TIN removed through anammox.
- AnAOB including *Candidatus_Kuenenia* and *Candidatus_Scalindua* were enriched in MSPDA-MBBR within 106 days.
- In MSPDA, thiosulfate promoted the electron provision rate and enhanced the nitrite accumulation.
- Oxidation of sulfide and thiosulfate to S^0^ and sulfate, respectively in autotrophic denitrification, boosted nitrite accumulation.
- The feasibility of SANIA process was fully demonstrated by 90%, 93%, and 61% removal efficiency of COD, ammonium, and TIN, respectively with the concentrations in effluent below 10, 3, and 13 mgN/L.
- Through combination of SANIA and organics capture (i.e., CEPT), it’s promising to achieve energy-neutral for wastewater treatment.

## Acknowledgements

This work was supported by the Hong Kong Innovation and Technology Commission (ITC-CNERC14EG03), the Research Grants Council of the Hong Kong SAR (T21-604/19-R), the International Science and Technology Cooperation Project of Huangpu District, Guang-zhou (Grant No. 2020GH04), Ghent University (BOF/STA/202109/022), Korea Ministry of Environment (2022003050003).

