## Supplemental information for "The oxygen-induced thiosulfate production during Sulfate reduction, Autotrophic denitrification, Nitrification and Anammox (SANIA) integrated process towards next-generation mainstream wastewater treatment"

**List of contents**

**SI 1:** Description of the reactor set-up and MBBR carriers of SANIA system

**SI 2:** Calculation

**SI 3:** MSPDA-MBBR performance data

**SI 4:** Performance of SANIA system

**SI 5:** Details for microbial analysis

**SI 6:** Critical values for energy calculation

**List of tables**

**Table S1** Characteristics of three types of carriers.

**Table S2** Performance of SPDA-MBBR in Period I during days 0-319.

**Table S3** Performance for completed SANIA system in Period II during days 325-455.

**Table S4** Diversity analysis of microbial samples.

**Table S5** Energy evaluation for the combination of chemical enhanced wastewater treatment and SANIA.

**List of figures**

**Fig. S1** Configuration of SANIA in (a) Period I and (b) Period II.

**Fig. S2** Profiles of microbial community in SPDA-MBBR at: (a) phylum level and (b) genus level during the Period I.

**Fig. S3** Profiles of microbial community in SANIA system on the end of the Period II (day 455) at: (a) phylum level and (b) genus level.

**SI 1: Description of the reactor set-up and MBBR carriers of SANIA system**

**1.1 Description of MBBR carriers**

K3 carries in anoxic MSPDA-MBBR were obtained from a lab-scale anoxic autotrophic sulfur-based denitrification reactor which has already been run for two years in our laboratory (Leung, 2019). Carriers in N-MBBR were taken from a lab-scale nitrification MBBR with over 200 days’ operation with influent ammonium as 100 mgN/L and effluent ammonium below 10 mgN/L. Detailed information for carriers was listed in Table S1.

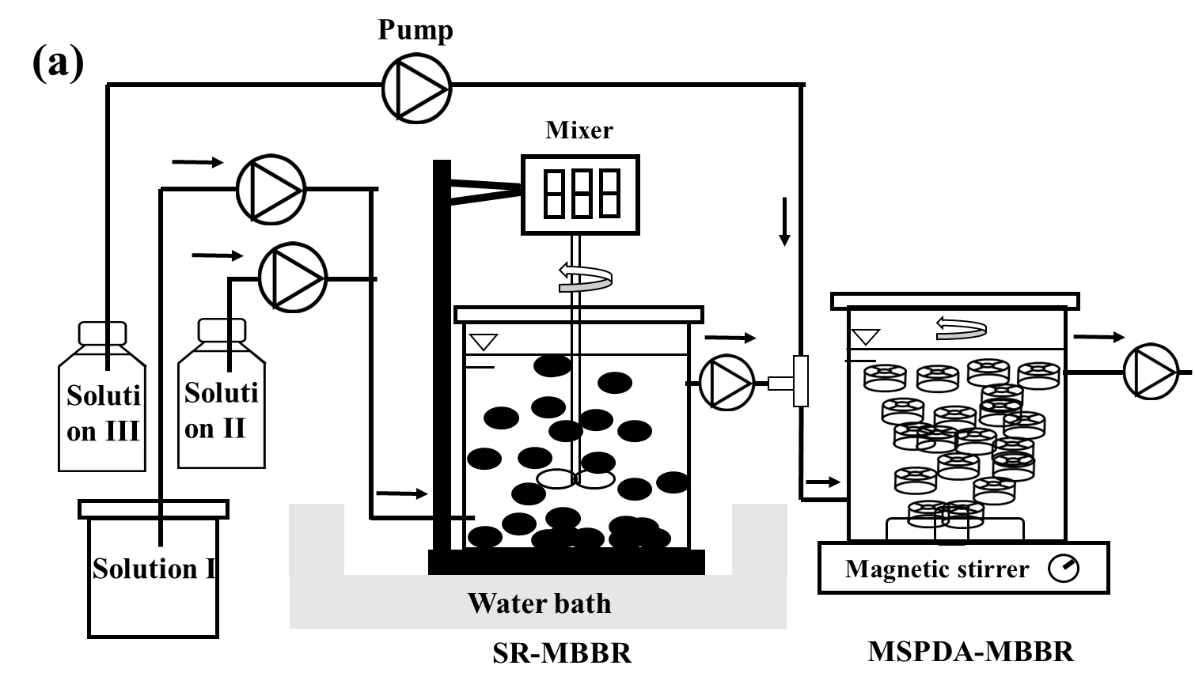

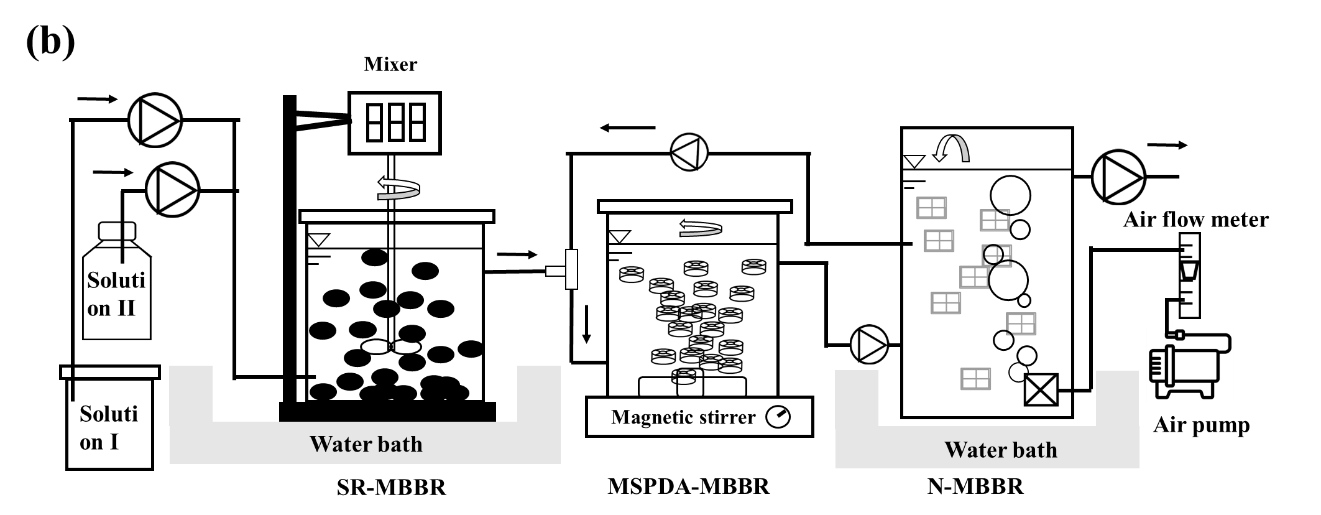

**Fig. S1** Configuration of SANIA in (a) Period I and (b) Period II.

**Table S1** Characteristics of three types of carriers.

| Type of Carrier | BioChip 30^TM^ | AnoxKaldnes^TM^ K3 | ActiveCell 920 |
| --- | --- | --- | --- |
| Depth (mm) | 1.1 | 10 | 12 |
| Diameter (mm) | 30 | 25 | 12 |
| Specific area (m^2^/m^3^) | 5,500 | 500 | 680 |
| Effective surface area (m^2^) | 2.2 | 0.25 | 1.22 |

**1.2 The composition of synthetic wastewater**

During the Period I, solution I and II were presented as reported for SR-MBBR (#refer#). Solution III (mimic nitrified effluent) was applied in MSPDA-MBBR. The Solution III is a synthetic solution with 650-700 mg NO_3_^-^-N/L. Flow rate of SR-MBBR effluent and Solution III were 25 mL/min and 1.3 mL/min respectively.

**SI 2: Calculation**

**2.1 Kinetic rate analysis for autotrophic denitrification and anammox**

Ammonium uptake by growth of bacteria was neglected because of the low yield coefficients of SOB and anammox bacteria (AnAOB) (Cui et al., 2019; Lotti et al., 2014). The surface-specific rates of autotrophic denitratation, denitritation and anammox in MSPDA-MBBR were determined as described:

for autotrophic denitrification,

r_d,_ _NO3_= -d[NO_3_^-^-N]/dt/dA

r_d,_ _NO2_= r_d,_ _NO3_ - d[NO_2_^-^-N] /dt/dA

for PD/A,

r_a,_ _NH4_ = -d[NH_4_^+^-N]/dt/dA

r_a,_ _NO2_= 1.32 r_a,_ _NH4_

r_d,_ _NO3_= -d[NO_3_^-^-N]/dt/dA+0.26r_a,_ _NH4_

r_d,_ _NO2_= r_d,_ _NO3_- r_a,_ _NO2_ - d[NO_2_^-^-N] /dt/dA

A is the specific surface area of the biofilm in MSPDA-MBBR (0.25 m^2^/L) r_d,_ _NO3_ and r_d,_ _NO2_ are the nitrate and nitrite utilization rates through denitratation and denitritation, r_a,_ _NH4_ and r_a,_ _NO2_ are the ammonium and nitrite utilization rates through Anammox (mg N/(m^2^·h)). 1.32r_a,_ _NH4_ and 0.26r_a,_ _NH4_ refer to NO_2_^-^-N consumption and NO_3_^-^-N production via anammox reaction. For batch tests, A is 0.1 m^2^/L. NO_2_^-^-N consumption and NO_3_^-^-N production via anammox reaction were determined to be 1.27 and 0.14, respectively in batch C.

**2.2 Calculation recycle ratio**

The recycle ratio X is calculated based on steady state operation (assume ammonium is completely oxidized in N-MBBR and nitrate in MSPDA-MBBR is removed via MSPDA with NO_3_^-^-N/NH_4_^+^-N ratio as 1.0)

Inf. NH_4_^+^-N = (Nae1 + X*Nae3)/(1+X) = (Nae1+2* Nae3)/3

Eff. NH_4_^+^-N = Nae2

Inf.TIN-N =2*Ntine3+ Ntine1)/3

Eff.TIN-N= Ntine2

In which, Ntine1, Ntine2 and Ntine3 refer to TIN concentration in SR-MBBR, MSPDA-MBBR and N-MBBR effluent, respectively.

Therefore, ammonium and nitrate in MSPDA-MBBR influent:

Nai2=Nae1/(1+X)

Nni2= XNne3/(1+X)

ammonium and nitrate in MSPDA-MBBR effluent:

Nne2=0

Nae2= Nae1/(1+X) - XNne3/(1+X)

ammonium and nitrate in effluent of N-MBBR

Nae3=0

Nne3=Nae2

=Nae1/(1+2X)

In which Nai2, Nae1, Nne2 and Nae3 represent ammonium concentrations in MSPDA-MBBR influent, in effluent of SR-MBBR, MSPDA-MBBR and N-MBBR, respectively. Nni2 and Nne3 refer to nitrate in MSPDA-MBBR and N-MBBR effluent. Ammonium was utilized for SRB’s growth in SR-MBBR was neglected, Nae1 is equal to ammonium in SR-MBBR influent which is 30 mgN/L. Thus, recycle ratio 2.0 is determined to achieve 6 mg/L NO_3_^-^-N in effluent (Nne3).

**2.3 Determination of the anammox and denitrification contributions to TIN removal**

Anammox and denitrification contribution to TN removal in MSPDA process were calculated. Calculations are based on the nitrogen concentration change from influent to effluent and stoichiometry reported by (Strous et al., 1998) as below and details after recycling were presented:

TIN removal via Anammox = [(Inf. NH_4_^+^-N - Eff. NH_4_^+^-N) + 1.32*(Inf. NH_4_^+^-N - Eff. NH_4_^+^-N) - 0.26*(Inf. NH_4_^+^-N - Eff. NH_4_^+^-N)]*100%/(Inf.TIN-N - Eff.TIN- N)

TIN removal via denitritation = (Inf.TIN-N - Eff.TIN-N) - (Inf. NH_4_^+^-N - Eff. NH_4_^+^-N) - 1.32*(Inf. NH_4_^+^-N - Eff. NH_4_^+^-N) + 0.26*(Inf. NH_4_^+^-N - Eff. NH_4_^+^-N)]*100%/(Inf.TIN-N - Eff.TIN-N)

After application of N-MBBR with recycle ratio as 2.0 in SANIA during Period II,

Inf. NH_4_^+^-N = (Nae1 + X*Nae3)/(1+X) = (Nae1+2* Nae3)/3

Eff. NH_4_^+^-N = Nae3

Inf.TIN-N =(2*Ntine3+ Ntine1)/3

Eff.TIN-N= Ntine2

In which, Ntine1, Ntine2 and Ntine3 refer to TIN concentration in SR-MBBR, MSPDA-MBBR and N-MBBR effluent, respectively.

**SI 3: MSPDA-MBBR performance**

**Table S2** Performance of MSPDA-MBBR in Period I.

| Parameters | Days 98–135 | | Days 217–240 | | Days 241–319 | |
| --- | --- | --- | --- | --- | --- | --- |
| Influent sulfate (mg S/L) | 29.3±7.3 | | 25.5±4.4 | | 25.3±1.8 | |
| Influent sulfide (mg S/L) | 12.1±3.0 | | 16.0±4.3 | | | |
| Influent thiosulfate (mg S/L) | 10.4±1.8 | | 3.9±2.4 | | 6.4±2.9 | |
| Effluent sulfate (mg S/L) | 43.5±7.6 | | 41.4±6.1 | | 41.1±2.8 | |
| Effluent sulfide (mg S/L) | 0.0 | | 0.0 | | 0.0 | |
| Effluent thiosulfate (mg S/L) | 0.0 | | 0.0 | | 0.0 | |
| Surface-specific sulfide removal rate (g S/(m^2^·d)) | | 1.7±0.4 | | 2.3±0.6 | | |
| Surface-specific thiosulfate removal rate (g S/(m^2^·d)) | | 1.5±0.3 | | 0.6±0.3 | | 0.9±0.4 |
| Sulfur balance (%) | 84 | | 83 | | 72 | |
| Influent organics (mg COD/L) | 35.4±18.2 | | 52.9±10.7 | | 46.2±6.3 | |
| Effluent organics (mg COD/L) | 23.9±13.9 | | 39.4±12.5 | | 34.7±5.7 | |
| Influent ammonium (mgN/L) | 29.2±2.9 | | 28.5±2.0 | | 29.9±2.1 | |
| Influent nitrate (mgN/L) | 29.3±4.1 | | 32.2±0.7 | | 34.7±2.2 | |
| Effluent ammonium (mgN/L) | 24.8±2.9 | | 21.1±2.9 | | 19.9±1.8 | |
| Effluent nitrate (mgN/L) | 10.6±2.3 | | 13.8±0.7 | | 14.4±1.5 | |
| Effluent nitrite (mg/L) | 6.8±2.1 | | 6.9±0.8 | | 5.8±1.1 | |
| Surface-specific ammonium removal rate (g N/(m^2^·d)) | 0.7±0.4 | | 1.3±0.4 | | 1.4±0.4 | |
| Surface-specific denitrification rate (g N/(m^2^·d)^b^) | 2.7±0.6 | | 2.7±0.1 | | 2.9±0.4 | |
| Surface-specific nitrite production rate (g N/(m^2^·d)) | 1.0±0.3 | | 1.0±0.1 | | 0.8±0.2 | |
| TIN removal via anammox (%) | 52 | | 77 | | 81 | |
| Biofilm attached total solids (g ATS/L) | 6.8±1.9 | | 8.0±2.7 | | 4.9±1.5 | |
| Biofilm attached volatile solids (g AVS/L) | 6.3±1.9 | | 7.7±2.6 | | 4.7±1.5 | |
| Effluent TSS (mg /L) | - | | - | | 17.6±2.7 | |
| Effluent VSS (mg/L) | - | | - | | 16.9±1.4 | |

Note: the data were obtained in 70-106d, 217-240d, and 271-319d; Volumetric denitrification rate (kg N/(m^3^ ·d)) = (Influent nitrate - Effluent nitrate) (mg N/L) х 24 (h/d) /HRT (h). b Surface-specific denitrification rate (mgN/(m^2^ ·h)) = (Influent nitrate - Effluent nitrate) (mg N/L) х reactor volume (L)/total carrier area (m^2^).

**SI 4: Performance of SANIA system**

**SI 5: Details for microbial analysis**

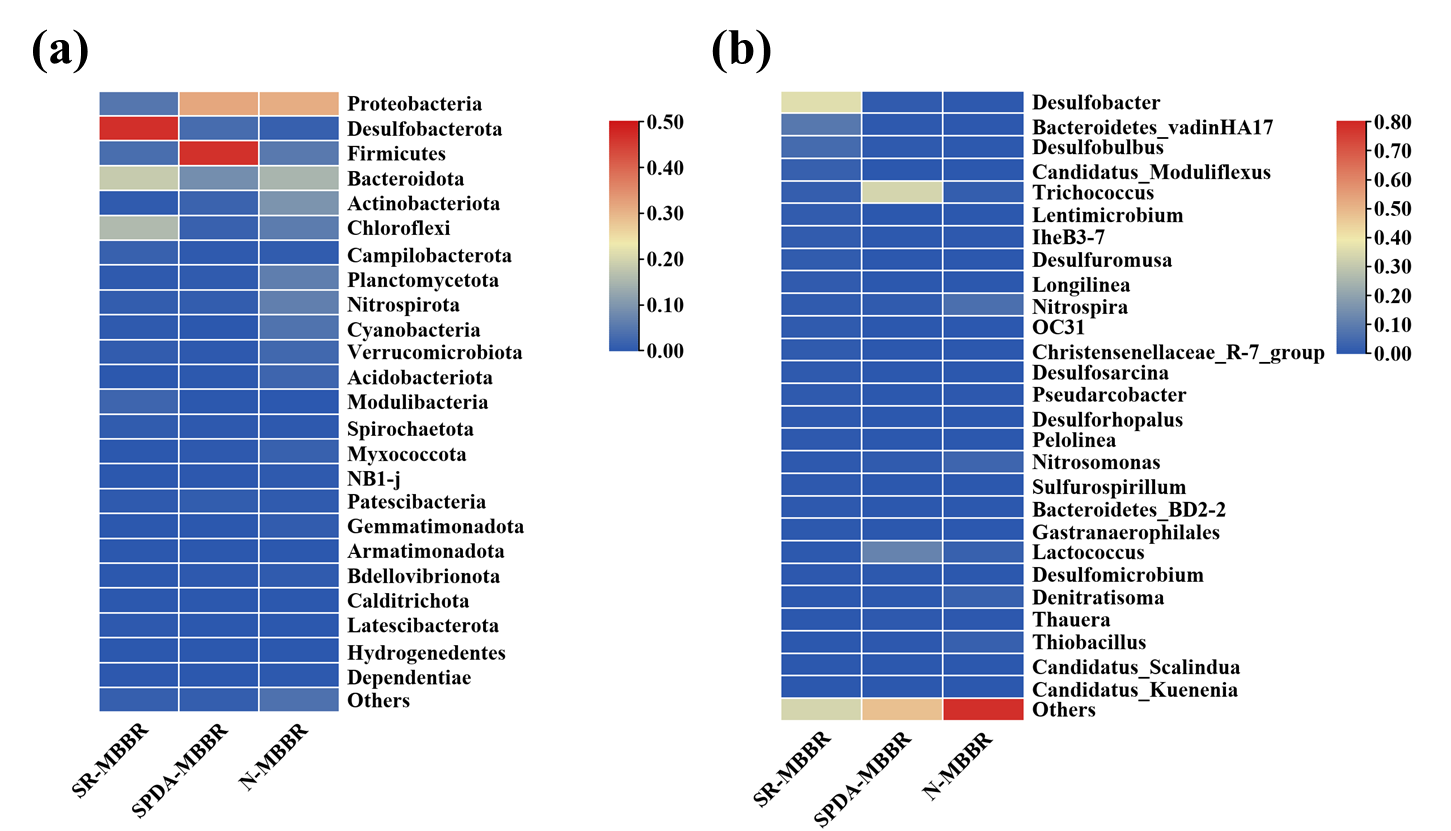

**Fig. S3** Profiles of microbial community in SANIA system on the end of the Period II (day 455) at: (a) phylum level and (b) genus level.

After connection of N-MBBR in complete SANIA system, main functional bacteria of sludge samples in genus level taken from SR-MBBR, MSPDA MBBR and N-MBBR on day 509 were shown as Fig. S4. As SANIA process operated for 73 days, SRB representative genus *Desulfobacter* and *Desulfobulbus* both increased to 36.2% and 5.1% separately in SR-MBBR. Fermenting bacteria like *Lactococcus* and *Trichococcus* were decreased to 1.8% and 0.3%, pgrespectively. At meanwhile in MSPDA-MBBR, total abundance of denitrifying bacteria decreased to 1.4% with dominant genus as *Denitratisoma* (0.4%), *Thauera* (0.5%) and *Thiobacillus* (0.2%). For anammox bacteria, total abundance including, *Candidatus Scalindua* (0.07%) and *Candidatus Kuenenia* (0.03%) were decreased with a total abundance of 0.1% (*Planctomycetes* decreased to 0.5%) with actual HRT decreased to 20 min. Decrease of AnAOB to denitrifiers ratio was also in accordance with less nitrogen removal via anammox (73.8%). In biofilm sample of N-MBBR, AOB and NOB were found to be R-strategists *Nitrosomonas* (3.7%) and a K-strategist *Nitrospira* (6.4%). *Nitrospira* was normally found dominant in partial nitrification process with operational DO lower than 1 mg/L in PN/A MBBR (Gilbert et al., 2014), in corresponding to limited DO (2.7±0.3 mg/L) applied in N-MBBR. At meanwhile, *Candidatus Scalindua* (0.04%), *Candidatus Kuenenia* (0.01%) and *Candidatus Brocadia* (0.05%) were also found in N-MBBR on day 455. As presented in Table S4, OTUs change indicated microbial diversity in SR-MBBR was increased in phase 2, decreased in phase 3 and further decreased after N-MBBR connection. In accordance with OTUs results, Index like Shannon, Simpon, Chao1 and ACE were increased first and then decreased. Different from SR-MBBR, OTUs in MSPDA-MBBR were maintained around 1100 while it largely increased to 1486 after N-MBBR connection, indicating nitrified effluent effect on MSPDA.

**Table S4** Diversity analysis of microbial samples.

| Sample | OTUs | Shannon | Simpson | Chao1 | ACE | Coverage |
| --- | --- | --- | --- | --- | --- | --- |
| MSPDA-MBBR. Day 106 | 1154 | 6.462 | 0.968 | 1400.392 | 1424.306 | 0.995 |
| MSPDA-MBBR. Day 240 | 1178 | 6.585 | 0.969 | 1439.462 | 1448.705 | 0.995 |
| MSPDA-MBBR. Day 319 | 1104 | 6.892 | 0.968 | 1167.182 | 1191.053 | 0.998 |
| SR-MBBR. Day 455 | 1040 | 5.099 | 0.869 | 1308.024 | 1327.360 | 0.995 |
| MSPDA-MBBR. Day 455 | 1486 | 5.442 | 0.869 | 1748.016 | 1779.530 | 0.994 |
| N-MBBR. Day 455 | 1103 | 7.679 | 0.984 | 1154.733 | 1150.578 | 0.998 |

Operational Taxonomic Units (OTUs) were classified with the sequence similarity over 0.97.

**SI 6: Critical values for energy calculation**

**Table S5** Energy evaluation for the combination of chemical enhanced wastewater treatment and SANIA.

| Parameters | Units | Value |
| --- | --- | --- |
| Influent COD | g/m^3^ wastewater | 500 (samples taken from Sai Kung Sewage Treatment Works, Hong Kong) |
| Influent NH_4_^+^-N | g/m^3^ wastewater | 30 (samples taken from Sai Kung Sewage Treatment Works, Hong Kong) |
| Energy contained in influent COD | kWh/m^3^ wastewater | 3.86 (McCarty et al., 2011) |
| Averagely energy consumption for conventional wastewater treatment | kWh/m^3^ wastewater | 0.45 (Wan et al., 2016) |
| The percentage of aeration energy for organics | % | 50 (Lotti, 2016) |
| The percentage of aeration energy for nitrogen | % | 25 (Lotti, 2016) |
| The percentage of aeration energy saving for nitrogen in SANIA | % | 20 |
| The percentage of pumping energy for sludge and wastewater | % | 6 (Sharma et al., 2011) |
| The percentage of mixing energy for MBBRs | % | 5 (Sharma et al., 2011) |
| The percentage of COD capture in CEPT | % | 80 (Ødegaard, 2016; Wan et al., 2016) |
| The conversion ratio of COD to methane | % | 40 (Wan et al., 2016) |
| The conversion ratio of methane to electricity | % | 40 (McCarty et al., 2011) |
| Water content for digested sludge | % | 95 |
| Energy consumption for sludge dewatering | kWh/m^3^ wastewater | 0.03 (Longo et al., 2016) |
| Electricity production from incineration | kWh/kg sludge | 0.01 (Abuşoğlu et al., 2017) |
